# Epigenomic and transcriptomic analyses reveal cnidocyte specialization in a sea anemone

**DOI:** 10.1101/2025.10.06.680633

**Authors:** Itamar Kozlovski, Adrian Jaimes-Becerra, Daria Aleshkina, Matan Levy, Yehu Moran

**Author notes:** Corresponding authors (IK); (YM).

## Abstract

Cnidarians, including corals, hydras, jellyfish, and sea anemones, possess specialized stinging cells called cnidocytes that function in prey capture and defense. These cells represent a striking evolutionary innovation and produce distinct types of organelles such as venom injecting nematocysts and mechanically entangling spirocysts. While their biomechanics and transcriptional regulation have been studied extensively, little is known about their epigenetic regulation. Here, we combined epigenetic profiling with RNA sequencing in the sea anemone *Nematostella vectensis* to explore regulatory programs underlying cnidocyte diversity. We identified cell-type-specific regulatory elements in promoter and enhancer regions and linked them to distinct gene expression programs. This analysis revealed fundamental differences between nematocytes and spirocytes and uncovered a previously unrecognized nematocyte population that expresses the Nep3 toxin but lacks most other toxins. These findings highlight the complexity of cnidocyte regulation and suggest greater cellular diversity within this defining cnidarian cell type than previously appreciated.

## Introduction

Cnidocytes, the explosive stinging cells unique to cnidarians, represent one of the most remarkable examples of cellular novelty in animal evolution ^1-3^. These cells, armed with specialized organelles called cnidae or cnidocysts, enable prey capture, defense, and various interspecific and intraspecific interactions ^4,5^. In sea anemones, these cells differentiate into distinct cell types, such as venom-injecting nematocytes and mechanically acting adhesive spirocytes used for entangling prey ^6^. A key feature of cnidocytes is their single-use nature: once discharged, they become nonfunctional and must be continuously replenished from progenitor pools ^7,8^. This combination of extreme specialization, diversity, and rapid turnover has positioned cnidocytes as an important system for studying the emergence and maintenance of novel cell types ^9^.

Classical studies elucidated the biomechanics of cnidocyte discharge and morphology ^6,10,11^, laying the foundation for understanding their biological function. More recently, molecular and single-cell transcriptomic approaches in *Nematostella vectensis* have revealed transcription factors and developmental trajectories that drive cnidocyte differentiation ^12-16^. Transcription factors such as *PaxA ^15^*, *Pou4 ^17^*, *ZNF845* ^18^, Jun and Fos ^14^, have been implicated in specifying cnidocyte fate and cell type identity. Moreover, miR-2022, the most conserved microRNA across Cnidaria, is expressed specifically in cnidocytes and regulate cnidae biogenesis ^19^. These studies have provided the first insights of the gene regulatory networks underlying cnidocyte development, but they capture only one layer of gene regulation.

In bilaterian systems, the interplay between transcription factors and chromatin architecture is central to cell fate specification ^20^. Histone modifications, such as H3K27ac, mark active promoters and enhancers, providing a regulatory landscape that defines lineage-specific gene expression ^21,22^. A growing body of evidence indicates that, in many contexts, chromatin accessibility and histone modifications define cell identity more robustly than RNA expression, with accessibility changes preceding transcription during lineage commitment ^23^ and histone or accessibility profiles alone sufficiently resolving cell identities and developmental trajectories that are often indistinguishable by RNA expression alone ^24-26^. By capturing regulatory states that reflect not only active but also repressed and poised genes, histone modifications offer critical insights into complex phenotypes such as neurogenesis ^27^ and regenerative capacity ^28^. Despite this well-established framework in other animals, the contribution of chromatin level regulation to cnidocyte biology remains largely unexplored. While genome wide assays of chromatin accessibility and histone modifications have been performed in cnidarians at whole organism resolution ^29,30^, no study has yet profiled histone modifications specifically in cnidocytes. This gap limits our understanding of how transcriptional programs are integrated with cis-regulatory mechanisms in the evolution of novel cell types.

Here, we present the first cell type specific profiling of histone modifications in cnidocytes of the sea anemone *Nematostella vectensis*, a well-established cnidarian model ^31-33^. By establishing CUT&Tag profiling of H3K27ac with transcriptomic data, we uncover the cis-regulatory architecture that supports cnidocyte gene expression and cell type diversification. To achieve cell type-specific resolution, we employed transgenic reporter lines in which fluorescent proteins were driven by cnidocyte type-specific promoters: Ncol3, a structural protein broadly expressed across cnidocytes ^14,34^, and Nep3, a nematocyte-specific toxin ^35^. These lines enabled the selective isolation of general vs. nematocyte restricted cnidocyte populations for epigenomic and transcriptomic analysis. This work provides a framework for linking transcriptional states to underlying enhancer and promoter dynamics, offering new insights into the molecular regulation of cnidocyte differentiation and the chromatin basis of novel cell type emergence in early-branching animals.

## Results

### Imaging flow cytometry reveals cell type-specific reporter expression

To investigate lineage specific gene expression and histone modifications we utilized two transgenic reporter lines: (1) *Ncol3::mOrange2*, which employs the broadly expressed *minicollagen Ncol3* promoter to drive ubiquitous mOrange2 expression across cnidocytes ^14^; and (2) *Nep3::mOrange2,* which uses the *Nep3* toxin promoter to drive nematocyte-specific expression of the mOrange2 transgene ^35^.

To obtain a comprehensive, quantitative assessment of reporter expression across cell morphologies, we first employed imaging flow cytometry using the ImageStream system ^36^. This approach enabled high-throughput statistical analysis of single-cell morphology and reporter signal distribution, providing a detailed characterization of the expression profiles of the two reporter lines. To this end, tentacles from adult individuals of both reporter lines were dissected, enzymatically dissociated, and analyzed by imaging flow cytometry. A total of 545 morphological features were quantified and statistically assessed to identify the defining characteristics of mOrange-positive cells in each line **(Figure 1A).** To ensure data validity, only focused events detected by both cameras were retained, and analysis was restricted to stringently gated, viable single cells that were properly centered in the image field **(Figure S1A).** This analysis confirmed the presence of spirocytes among Ncol3⁺ cells but not in Nep3⁺ cells, and identified circular precursor nematoblasts expressing mOrange in both lines **(Figure 1B)**. Consistent with the absence of spirocytes, Nep3⁺ cells exhibited a significantly more circular morphology **(Figure 1C)**, reduced elongation **(Figure 1D)**, higher aspect ratio **(Figure 1E)**, and larger cytoplasmic area **(Figure 1F)**. To quantify differences between the two cell populations in an unbiased manner, cells were clustered and visualized using uniform manifold approximation and projection (UMAP). This analysis resolved four distinct clusters corresponding to spirocytes, nematocytes, large circular cells, and small circular cells **(Figure 1G)**. Nep3⁺ cells were enriched for circular cells and contained a smaller fraction of mature cnidocytes compared to Ncol3⁺ cells **(Figure 1H)**.

**Figure 1.**
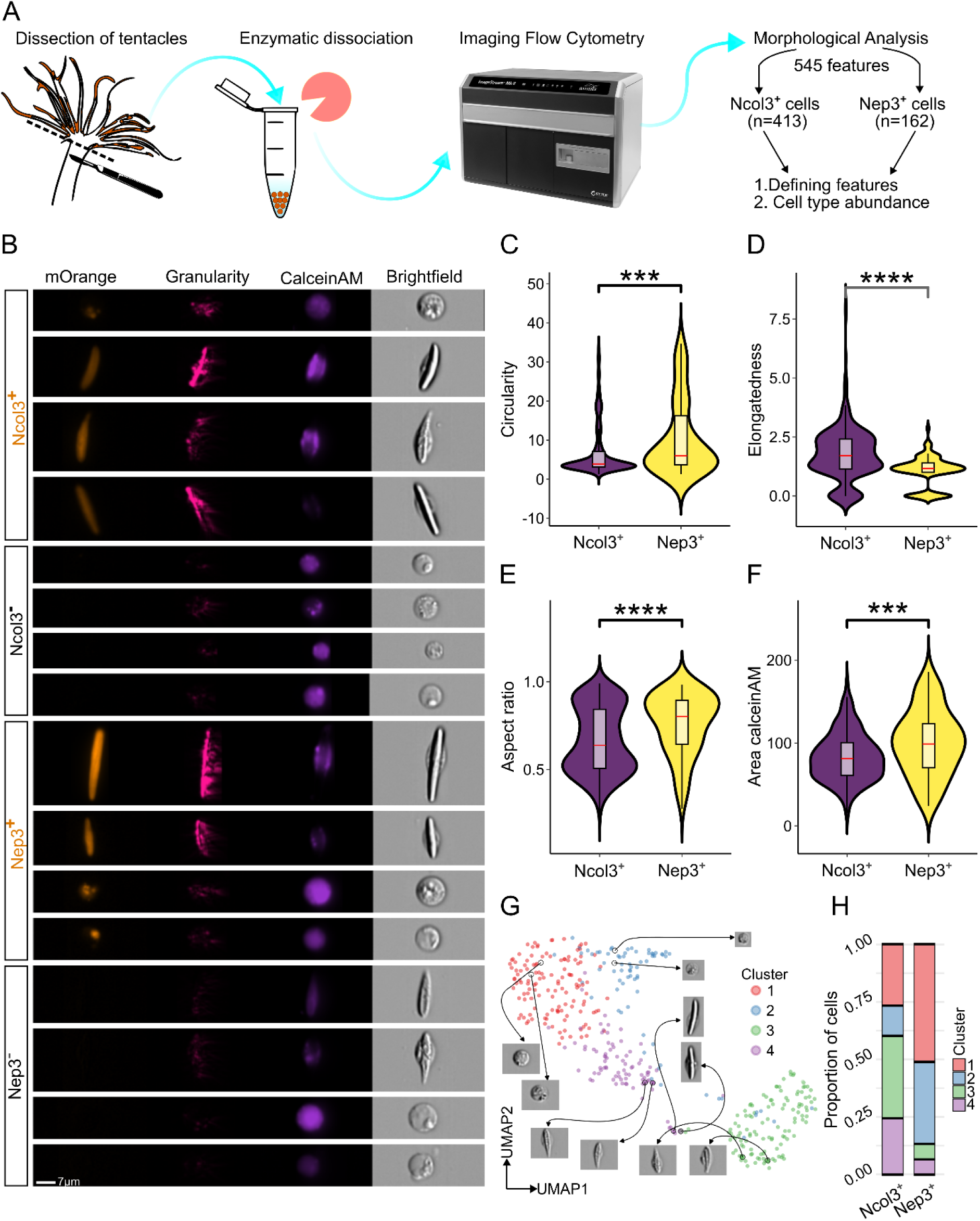
Cell type-specific expression patterns of Ncol3 and Nep3 reporter lines revealed by imaging flow cytometry. **(A)** Workflow for imaging flow cytometry of *Nematostella vectensis* reporter lines. Tentacles were dissected, enzymatically dissociated, and analyzed on the Amnis ImageStreamX platform. Morphological analysis of 545 extracted features was used to define distinguishing parameters and quantify cell-type abundance across Ncol3⁺ (n = 413) and Nep3⁺ (n = 162) cells. **(B)** Representative images of *Ncol3::mOrange* and *Nep3::mOrange* positive and negative cells showing mOrange fluorescence, intracellular granularity, CalceinAM staining, and brightfield morphology. Scale bar, 7 µm. **(C-F)** Quantitative comparison of key morphological parameters reveals significant differences between Ncol3⁺ and Nep3⁺ cells, including circularity (C), elongation (D), aspect ratio (E), and area circularity index (F). Two-sided t test was performed: ***p < 0.001; ****p < 0.0001. **(G)** UMAP projection of single-cell morphological features identifies four major clusters, with representative cell images highlighting distinct morphologies. **(H)** Distribution of Ncol3⁺ and Nep3⁺ cells across the identified clusters.

These quantitative findings were consistent with fluorescence microscopy of adult animals from the two reporter lines. Ncol3::mOrange2 individuals exhibited ubiquitous mOrange2 expression throughout the tentacles, whereas Nep3::mOrange2 individuals showed a more restricted pattern, with expression primarily concentrated at the tentacle tips **(Figure S1B)**. High-magnification fluorescent imaging of the reporter lines revealed distinct expression profiles between *Ncol3::mOrange* and *Nep3::mOrange*. In the *Ncol3::mOrange* line, fluorescence was observed across diverse cnidocyte morphologies, including elongated and smaller spiral cells, indicating broad expression among cnidocytes. In contrast, the *Nep3::mOrange* reporter was restricted to a narrower subset, with signal predominantly in elongated rod shaped cells, consistent with cell type-specific expression likely confined to nematocytes **(Figure S1C)**.

Together, these results demonstrate that Ncol3 functions as a general cnidocyte marker, whereas Nep3 is restricted to nematocytes. Both reporters are expressed in nematoblasts prior to their differentiation into mature cnidocytes.

### Transcriptomic profiling defines molecular signatures of Ncol3⁺ and Nep3⁺ cells

To complement the morphological characterization of Ncol3⁺ and Nep3⁺ reporter-expressing cells, we next performed transcriptomic profiling to define their underlying molecular signatures. Fluorescence-activated cell sorting (FACS) was used to isolate mOrange2-positive and negative fractions from both reporter lines, followed by RNA sequencing **(Figures 2A and S2A)**. This approach enabled a direct comparison of gene expression programs between general cnidocytes (Ncol3⁺), nematocytes (Nep3⁺), and their corresponding negative populations. By integrating these datasets, we aimed to identify cell type-specific transcriptional signatures and regulatory pathways that distinguish nematocytes from other cnidocyte cell types.

**Figure 2.**
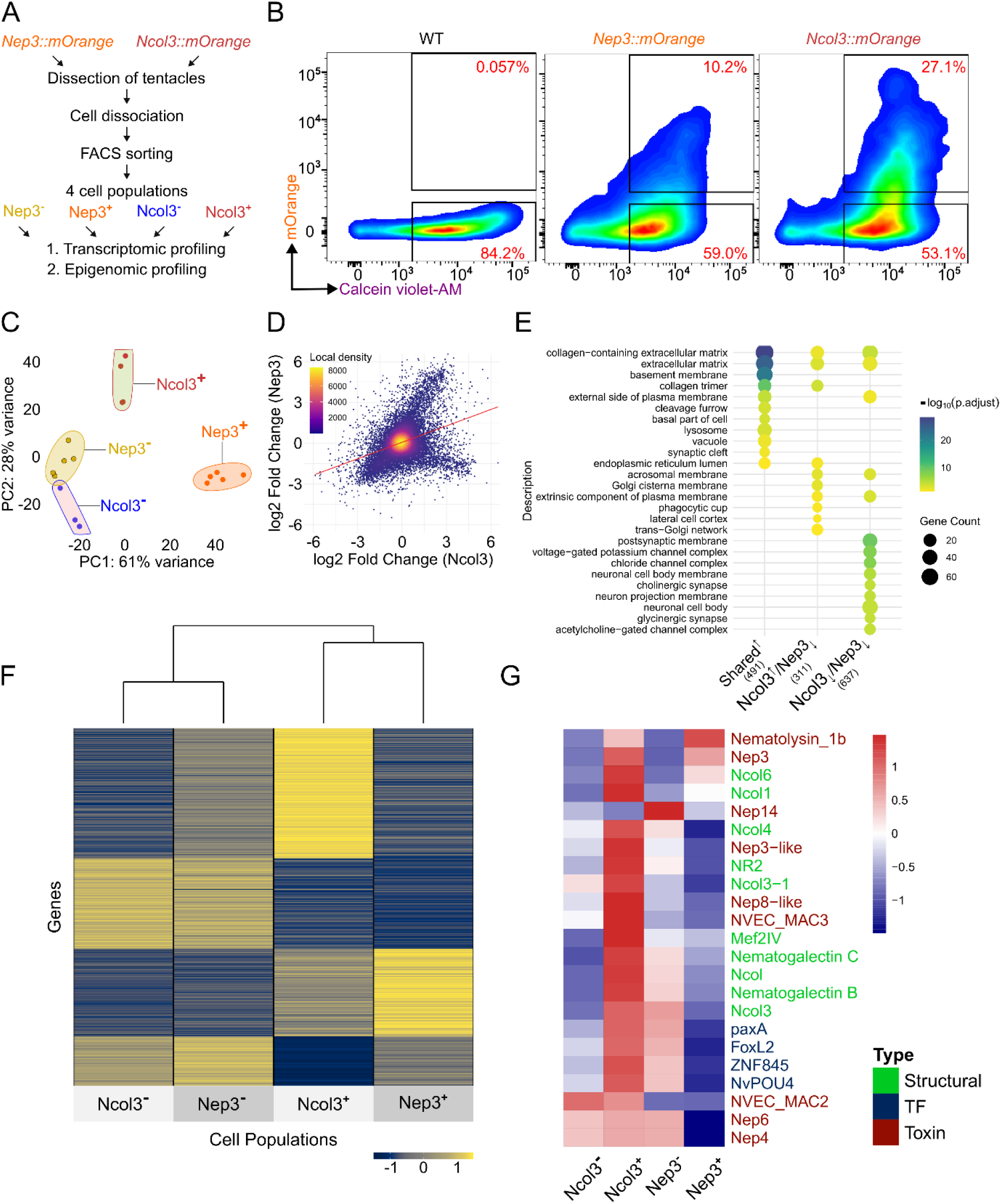
Transcriptomic profiling of FACS-sorted Ncol3⁺ and Nep3⁺ cnidocytes defines cell-type-specific programs. **(A)** Experimental workflow. Tentacles from transgenic *Nematostella vectensis* polyps expressing *Ncol3::mOrange* or *Nep3::mOrange* reporters were dissociated, and reporter-positive and negative populations were isolated by FACS for transcriptomic and epigenomic profiling. **(B)** Representative FACS plots showing gating strategy for wild-type, *Nep3::mOrange*, and *Ncol3::mOrange* polyps derived cells. **(C)** Principal component analysis of RNA-seq profiles separates the four sorted populations (Ncol3⁺, Ncol3⁻, Nep3⁺, Nep3⁻), with Ncol3⁺ and Nep3⁺ clustering apart from their respective negatives. **(D)** Scatter plot comparing differential expression between Ncol3⁺ and Nep3⁺ populations. Each dot represents a gene, colored by local density. **(E)** Gene Ontology enrichment of differentially expressed genes highlights distinct biological processes enriched in Ncol3⁺ and Nep3⁺ cells. Dot size indicates the number of genes, and color represents adjusted significance. **(F)** Heatmap of differentially expressed genes across the four populations reveals distinct expression signatures of Ncol3⁺ and Nep3⁺ cells. **(G)** Normalized expression heatmap of selected cnidocyte-related genes, including structural proteins (green), transcription factors (blue), and toxins (red).

In agreement with the imaging results **(Figures 1B and S1B)**, flow cytometry analysis confirmed that Ncol3⁺ cells were more abundant than Nep3⁺ cells, comprising ∼28% and ∼13% of the total population, respectively **(Figures 2B and S2B)**. Sorting validity was supported by marker expression: *mOrange2* was enriched in both positive fractions (log₂FC = 1.78, *padj* = 5.7 × 10⁻³ in Ncol3⁺; log₂FC = 3.04, *padj* = 1.6 × 10⁻¹⁰ in Nep3⁺), *Ncol3* was specific to Ncol3⁺ (log₂FC = 2.16, *padj* = 7.5 × 10⁻¹⁸), and *Nep3* was enriched in Nep3⁺ (log₂FC = 2.47, *padj* = 8.7 × 10⁻⁶). Principal component analysis (PCA) revealed distinct transcriptomic profiles associated with each cell population **(Figure 2C).** Correlation analysis of differential gene expression between Ncol3⁺ and Nep3⁺ cells revealed an overall positive relationship across the transcriptomes **(Figure 2D)**. Most genes clustered around the origin, reflecting similarly low expression in both populations, whereas subsets deviated along each axis, indicating cell type-specific regulation. The density distribution showed a greater proportion of genes upregulated in Ncol3⁺ cells but not in Nep3⁺ cells, consistent with the broader cell-type composition observed for Ncol3. These anti-correlated genes likely represent spirocyte-specific transcripts.

Gene ontology enrichment of molecular functions further supported the differential roles of the reporter-defined populations **(Figures 2E and S2C)**. Genes upregulated in both Ncol3⁺ and Nep3⁺ cells were enriched for structural and binding functions, including extracellular matrix structural constituents, collagen binding, integrin binding, and peptidase inhibitor activity, consistent with nematocyst structure and function. Ncol3⁺ specific genes showed enrichment for intracellular trafficking and enzymatic activities, such as glycosyltransferase activity, consistent with its broader role across cnidocyte types. In contrast, downregulated genes were strongly associated with neurotransmission-related functions, including ion channel activity, neurotransmitter receptor activity, and synaptic signaling, paralleling the cellular component enrichments observed in **Figure 2E**. Together, these results indicate that Ncol3⁺ and Nep3⁺ cells share core cnidocyte functions but diverge in cell type-specific structural and signaling capacities.

Unsupervised hierarchical clustering of differentially expressed genes across the four sorted populations (Ncol3⁻, Nep3⁻, Ncol3⁺, and Nep3⁺) revealed clear segregation between positive and negative fractions **(Figure 2F)**. Both Ncol3⁺ and Nep3⁺ cells clustered together and showed broad upregulation of cnidocyte-associated genes relative to their negative counterparts. Within the positive fractions, Ncol3⁺ cells exhibited wider gene activation, whereas Nep3⁺ cells displayed a more restricted profile, consistent with nematocyte specificity. Notably, a distinct set of genes was shared by Ncol3⁺ and Nep3⁻ populations, which include spirocytes, pointing to this cluster as a putative spirocyte-specific gene set. Indeed, re-analysis of a published single cell atlas ^12^ showed that the top anti-correlated genes (ranked by log₂ fold change; up in Ncol3⁺ and down in Nep3⁺) are highly expressed in spirocyte-annotated clusters **(Figure S2D)**.

Previously, Nep3 expression was detected in a subset of nematocytes, while other nematocytes lacked this toxin ^35^, suggesting heterogeneity in toxin composition. However, the precise toxin repertoire, structural proteins, and transcription factors of different nematocyte subtypes remain largely unexplored. To address this, we examined the expression of known toxins, structural proteins, and transcription factors across the four cell populations **(Figure 2G)**. We found that Nep3⁺ cells lacked most other toxins, with the exception of Nep3 itself and Nematolysin 1b. In contrast, Ncol3⁺ cells strongly expressed a range of structural proteins, including Ncol4, NR2, Ncol3, and Nematogalectin B and C, whereas Nep3⁺ cells were restricted to high expression of Ncol6. Strikingly, transcription factors implicated in nematogenesis such as PaxA, FoxL2, ZNF845, and NvPOU4, were all restricted to Ncol3⁺ cells and absent from Nep3⁺ cells. Together, these findings suggest that Nep3⁺ cells either represent a distinct, previously uncharacterized nematocyte subtype, or alternatively correspond to a specific cell state in which the transcription of most other toxins has not yet occurred.

### Establishing CUT&Tag Profiling in *Nematostella*

To define the cis-regulatory DNA landscape of cnidocytes, we established CUT&Tag profiling ^37^ of H3K27ac in *Nematostella vectensis*. While this method has previously been applied in *Hydra* ^30^, it has not been used in a cell type-specific context. To this end, we utilized the *Ncol3::mOrange* and *Nep3::mOrange* transgenic reporter lines, isolated positive and negative fractions by FACS, and subjected them to CUT&Tag sequencing **(Figure 3A).**

**Figure 3.**
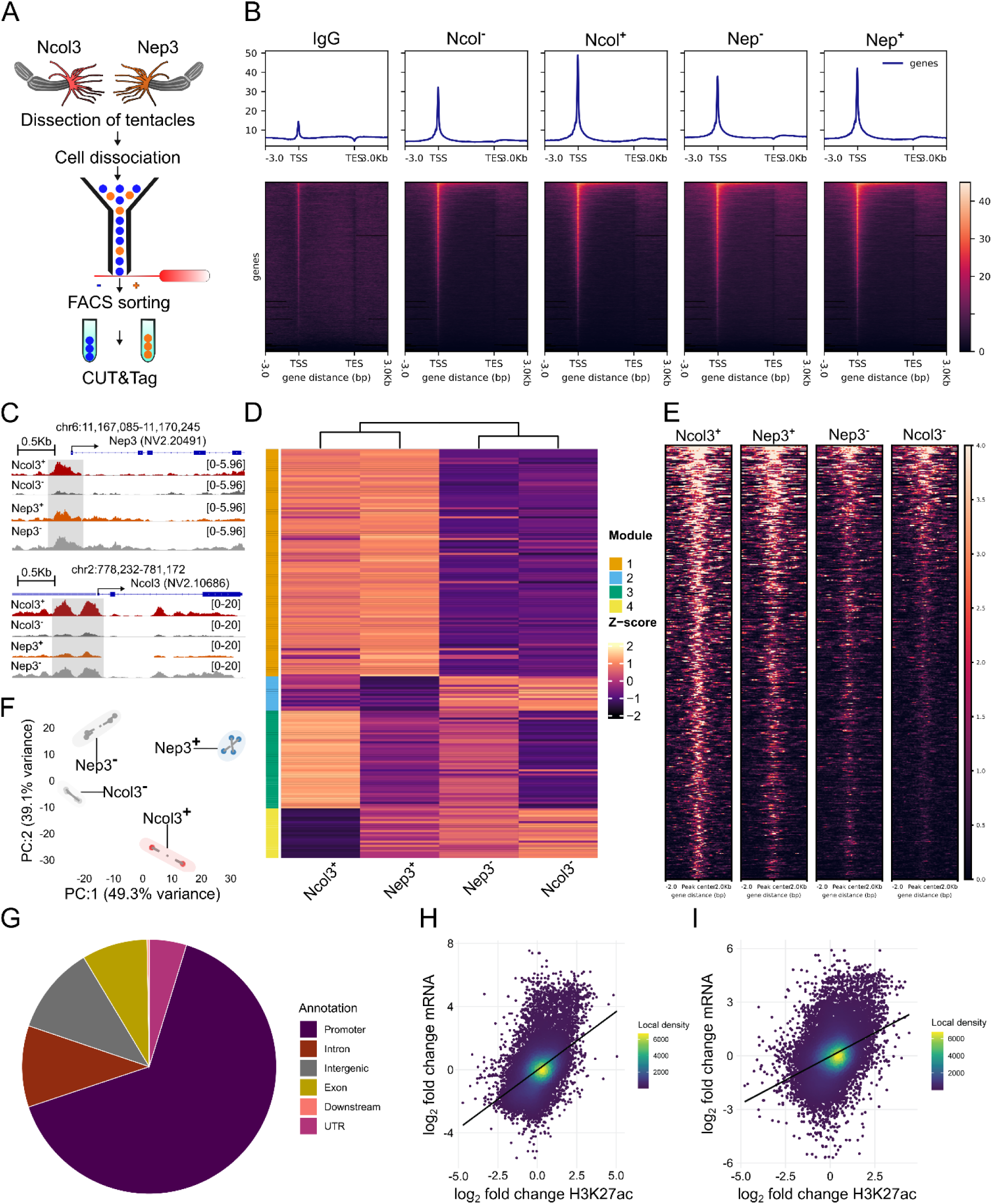
CUT&Tag profiling of H3K27ac reveals cell-type-specific regulatory landscapes in cnidocytes. (A) Experimental workflow. Tentacles from *Ncol3::mOrange* and *Nep3::mOrange* reporter lines were dissociated, FACS-sorted into positive and negative populations, and subjected to CUT&Tag for H3K27ac profiling. **(B)** Aggregate (profile) plots and heatmaps of normalized H3K27ac CUT&Tag signal around TSSs and TESs, averaged across all annotated genes. The y-axis shows mean normalized read density across four biological replicates (IgG, n = 2). **(C)** Genome browser tracks illustrating representative loci with cell type-specific H3K27ac enrichment at Ncol3 and Nep3 genes. **(D)** Hierarchical clustering of the top differential H3K27ac peaks in Ncol^+^ and Nep3^+^ cells (n = 189) identifies four modules of regulatory elements with distinct cell-type-specific patterns. **(E)** Heatmaps of H3K27ac signal intensities across Ncol3⁺, Ncol3⁻, Nep3⁺, and Nep3⁻ cells highlight module-specific enhancer activity. **(F)** Principal component analysis of H3K27ac profiles separates reporter-positive from negative populations, with clear distinction between Ncol3⁺ and Nep3⁺ cells. **(G)** Genomic distribution of H3K27ac peaks. **(H-I)** Correlation between H3K27ac enrichment and RNA expression in Nep3⁺ (H) and Ncol3⁺ (I) cells.

A series of quality control analyses confirmed that the CUT&Tag libraries were of high quality **(Figure S3)**. TapeStation profiles revealed the expected nucleosomal ladder with clear mono-, di-, and tri-nucleosome peaks **(Figure S3A)**. Sequencing depth and alignment rates were high and consistent across samples **(Figure S3B)**. Fragment length distributions of mapped reads further confirmed nucleosomal periodicity across all conditions, whereas IgG controls displayed an over-representation of sub-nucleosomal fragments **(Figure 3C)**. Duplication rates were generally low in biological samples but markedly elevated in IgG controls, consistent with the increased PCR amplification (18 vs. 14 cycles) required to recover sufficient DNA **(Figure S3D)**.

All samples exceeded the commonly used fraction of reads in peaks (FRiP) score (>0.15; **Figure S3E**), consistent with ChIP-seq guidelines and practices of the ENCODE project ^38^. Although IgG controls displayed elevated peak counts and higher FRiP values **(Figures S3E and S3F),** their peak annotations were broadly and non-specifically distributed across genomic features. By contrast, biological samples showed the expected enrichment at promoters as well as intronic and intergenic regions **(Figure S3G).** Importantly, a global analysis of CUT&Tag signal across all annotated genes revealed strong enrichment at transcription start sites (TSS) in biological samples but not in IgG controls **(Figure 3B)**, underscoring the specificity and overall high quality of the datasets for downstream analyses.

Inspection of the Ncol3 and Nep3 promoter regions revealed clear cell type-specific enrichment of CUT&Tag signal: the Nep3 promoter showed negligible signal in Ncol3⁻ cells, which are devoid of cnidocytes, but strong enrichment in Nep3⁺ cells, whereas the Ncol3 promoter was highly enriched in Ncol3⁺ cells and low in Ncol3⁻ cells **(Figure 3C)**. Building on these locus-specific observations, hierarchical clustering of the top 100 peaks from Nep3⁺ and Ncol3⁺ cells (189 unique peaks in total) revealed four distinct modules, separating positive from negative fractions and distinguishing shared vs. cell type-specific regulation **(Figure 3D)**. In line with the transcriptomic results, one of these modules was shared between Ncol3⁺ and Nep3⁻ cells, likely representing putative spirocyte-specific regulatory elements.

To extend this analysis, heatmap profiling of the top 500 differential H3K27ac peaks between Ncol3⁺ and Ncol3⁻ cells **(Figure 3E)** revealed strong enrichment in both Ncol3⁺ and Nep3⁺ fractions, with signal diminished in Nep3⁻ cells and further reduced in Ncol3⁻ cells, supporting the presence of both shared and cell type-specific regulatory activity in cnidocytes. Principal component analysis of CUT&Tag profiles **(Figure 3F)** corroborated these findings, showing clear separation of Ncol3⁺ and Nep3⁺ cells from their respective negative populations, while also distinguishing the two positive fractions from each other.

To gain further insight into the regulatory architecture of these differential peaks, annotation analysis showed that the majority (∼60%) localized to promoter regions, with additional peaks mapping to intronic (∼15%), intergenic (∼12%), exonic (∼7%), and UTR/downstream (∼6%) elements **(Figure 3G)**. Integration with RNA-seq data revealed a strong positive correlation between differential H3K27ac signal at promoters (< 3Kb from TSS) and gene expression changes, when restricted to differentially expressed genes (p < 0.05), in both Nep3⁺ (Pearson R = 0.81) and Ncol3⁺ (Pearson R = 0.74) cells **(Figs. 3H-I)**. This indicates that chromatin acetylation dynamics are tightly coupled to transcriptional activation in cnidocyte types. Collectively, these results confirm that our CUT&Tag datasets are robust and reproducible, enabling reliable discrimination of distinct cnidocyte identities.

### Functional and regulatory characterization of differential H3K27ac peaks

To further dissect the regulatory programs of cnidocyte types, we performed differential peak analysis and identified 13,628 differentially enriched regions out of 72,101 peaks across Ncol3⁺ vs. Ncol3⁻, Nep3⁺ vs. Nep3⁻, and Ncol3⁺ vs. Nep3⁺ comparisons. Differential H3K27ac peaks revealed clear cell type-specific activity **(Figure 4A)**, with Ncol3⁺ up peaks enriched in Ncol3⁺ but not Ncol3^-^ cells, and Nep3⁺ up peaks enriched exclusively in Nep3⁺ cells. Distance-to-TSS distributions showed a modest but significant shift (KS test: D = 0.03, p = 0.049), with ∼60% of Ncol3⁺-enriched peaks located within 1-3 kb of TSSs vs. ∼55% for Nep3⁺ peaks, while Nep3⁺ peaks were relatively more distal (>5 kb) **(Figure S4A)**. Overlap analysis revealed that Ncol3⁺ (3,815 peaks) and Nep3⁺ (4,787 peaks) share 2,276 regions, corresponding to ∼60% of Ncol3⁺ and ∼48% of Nep3⁺ peaks **(Figure S4B)**. This overlap was highly significant (Fisher’s exact test: odds ratio = 38.7, p < 2.2 × 10⁻¹⁶), indicating a core set of shared regulatory elements alongside large fractions of cell type-specific enhancers. Together, these results demonstrate that cnidocytes are defined by both shared and distinct chromatin landscapes that underpin their divergent transcriptional programs.

**Figure 4.**
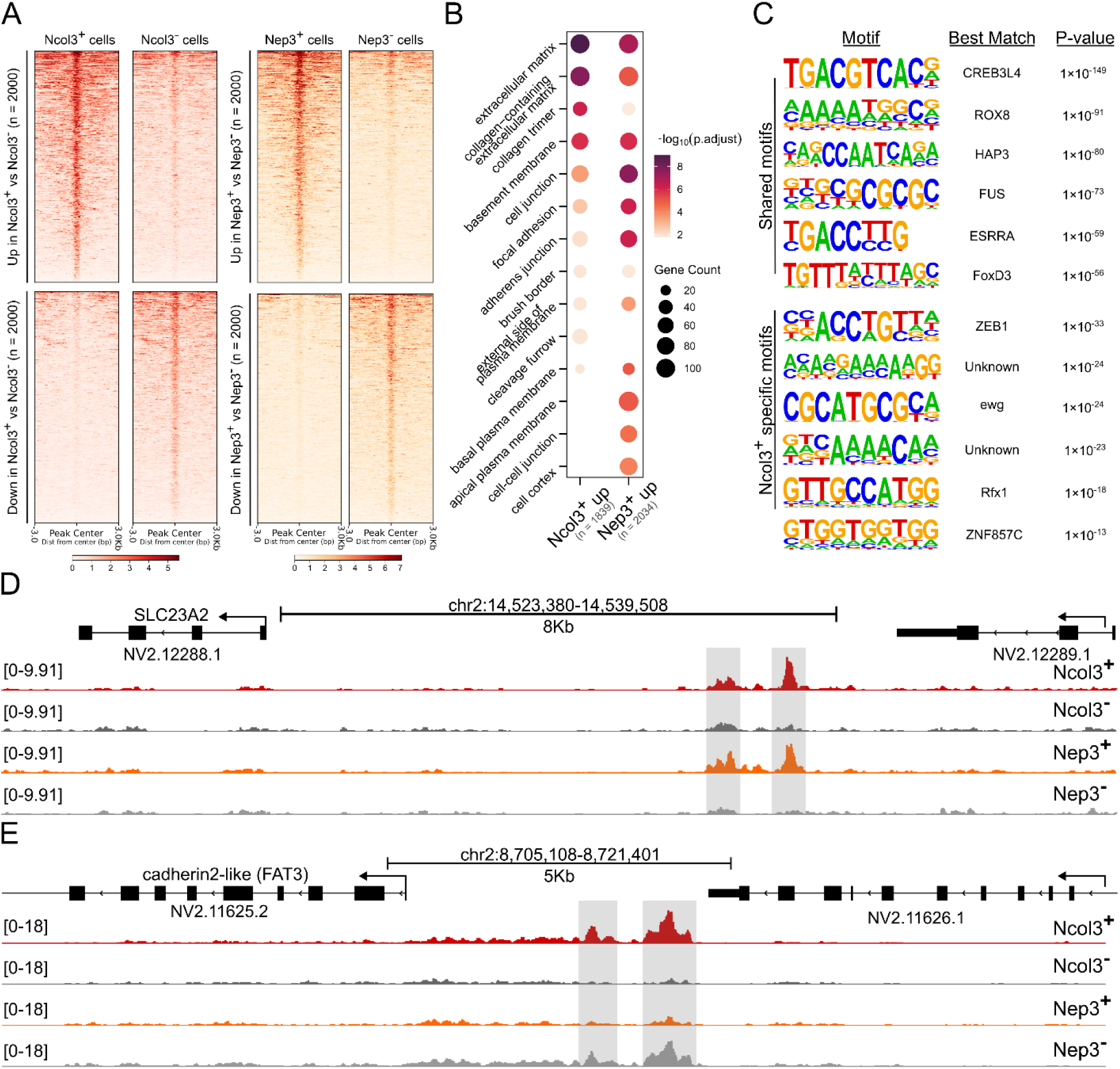
Analysis of H3K27ac peaks reveals shared and cell type-specific regulatory programs in cnidocytes. **(A)** Heatmaps of normalized H3K27ac CUT&Tag signal at the top 2,000 differentially enriched peaks across Ncol3⁺, Ncol3⁻, Nep3⁺, and Nep3⁻ populations. Separate panels show peaks upregulated in Ncol3⁺ vs. Nep3⁺ cells (top) and downregulated peaks (bottom). **(B)** Gene Ontology enrichment of genes associated with differentially acetylated regions. Dot size indicates the number of genes, and color denotes statistical significance (adjusted p-value). **(C)** Motif enrichment analysis identifies shared transcription factor motifs (top) and Ncol3⁺ specific motifs (bottom). Motifs are ranked by statistical significance, with predicted matches to known transcription factors shown. **(D)** Genome browser tracks illustrating a putative enhancer ∼7 kb upstream of the sodium-dependent L-ascorbate transporter gene (SLC23A2). This region is enriched in both Ncol3⁺ and Nep3⁺ cells but absent in negative populations. **(E)** Genome browser tracks of a putative enhancer ∼4 kb upstream of the cadherin2-like (FAT3) gene. This enhancer is enriched in Ncol3⁺ and Nep3^-^ cells and absent in Nep3⁺ cells, consistent with spirocyte-specific regulation.

Consistent with these chromatin-level distinctions, Gene Ontology (GO) enrichment of genes containing upregulated H3K27ac peaks in their promoters highlighted both cellular components and molecular activity differences between the two cnidocyte cell types. At the cellular component level **(Figure 4B)**, Ncol3⁺-associated genes were strongly enriched for extracellular matrix related categories such as *extracellular matrix*, *collagen-containing extracellular matrix*, and *collagen trimer*, reflecting their structural role in cnidocyte scaffolding. By contrast, Nep3⁺ associated genes were enriched for plasma membrane and junction-associated terms including *cell junction*, *adherens junction*, *apical plasma membrane*, and *cell cortex*, consistent with nematocytes structure. At the molecular function level **(Figure S4C)**, Ncol3⁺ promoter-associated genes were significantly enriched for *collagen binding* and *extracellular matrix structural constituent* activity, whereas Nep3⁺ promoter-associated genes showed preferential enrichment for categories such as *actin binding*, *monocarboxylic acid transmembrane transporter activity*, and *protein tyrosine kinase activity*. Both populations also shared enrichment in transcriptional regulatory functions (e.g., *sequence-specific DNA binding* and *transcription factor activity*), underscoring common regulatory requirements while revealing distinct cell type-specific molecular specializations.

Motif enrichment analysis of upregulated H3K27ac peaks revealed both shared and cell type-specific transcription factor binding motifs. Among the shared motifs, we identified highly significant matches to transcriptional regulators including CREB3L4, ROX8, HAP3, FUS, ESRRA, and FoxD3 (p-values ranging from 1×10⁻¹⁴⁹ to 1×10⁻⁵⁶), suggesting common regulatory inputs across cnidocyte types. In contrast, Ncol3⁺-specific peaks were enriched for motifs corresponding to ZEB1, ewg, Rfx1, and ZNF857C, as well as several uncharacterized motifs, pointing to distinct transcriptional regulators associated with the structural and functional specialization of Ncol3⁺ cells, which also contains spirocytes. Together, these results indicate that while Ncol3⁺ and Nep3⁺ cells share a core set of regulatory motifs, Ncol3⁺ cells also engage additional factors that may underlie their cell types-specific promoter and enhancer activities. Consistently, analysis of putative enhancers located >3 kb from TSSs **(Figure S4D)** revealed a large block of shared activity alongside distinct Ncol3⁺ and Nep3⁺ specific clusters. This pattern suggests that common transcription factors coordinate the shared enhancer modules, whereas Ncol3⁺ biased motifs drive cell type-specific enhancer activity that is absent in Nep3⁺ cells. For example, we identified a putative enhancer ∼7 kb upstream of the SLC23A2 gene, which encodes a sodium-dependent L-ascorbate transmembrane transporter **(Figure 4D)**. This enhancer was strongly enriched in both Ncol3⁺ and Nep3⁺ cells but nearly absent in their negative counterparts, and the corresponding transcript was significantly upregulated in Nep3⁺ cells (log₂FC = 5.25, padj = 3.8 × 10⁻⁷). Conversely, a putative enhancer located ∼4 kb upstream of the cadherin2-like (FAT3) gene **(Figure 4E)** was specifically enriched in Ncol3⁺ and Nep3⁻ cells, likely representing a spirocyte-associated regulatory element. Consistent with this pattern, FAT3 expression was upregulated in Ncol3⁺ cells and downregulated in Nep3⁺ cells (log₂FC = 2.14 and -1.18, padj = 2.53 × 10⁻⁸ and 2.78 × 10⁻⁴, respectively).

### Differential peak analysis reveals cell type specific regulation of transcription factors

To investigate the regulatory programs underlying cell type specificity, we focused on proximal promoter H3K27ac peaks associated with transcription factor (TF) genes. Out of the significant ∼24,000 peaks quantified, 803 overlapped with promoters (<3 kb from TSS) of TFs identified in our annotation (600 unique TF genes). Pairwise comparisons of Ncol3⁺ and Nep3⁺ vs. their negative counterparts highlighted dozens of transcription factors with cell type restricted promoter activity **(Figure 5A)**. In line with this, hierarchical clustering **(Figure S5A)** grouped candidate regulators according to their distinct acetylation patterns. Examples include zinc finger and homeobox transcription factors such as ZNF333, GBX2, SCRT2, and PTF1A, each showing localized acetylation peaks in Ncol3⁺, both Ncol3⁺ and Nep3⁺ cells, or their negative counterparts **(Figures S5B-E)**. Previously described regulators such as Cnido-Jun, Cnido-Fos ^14^ and the zinc finger protein ZNF845 ^18^ were, indeed, highly acetylated in a nematocyte specific manner **(Figures 5C-E)**. Notably, this analysis uncovered a set of previously undescribed transcription factors that exhibited nematocyte-specific acetylation, including homologs of FoxC, TCF15, and GLI4 **(Figures 5F-H)**. Consistent with the CUT&Tag results, these factors were also significantly upregulated in our RNA-seq data: FoxC (log₂FC = 3.06 in Ncol3⁺ and 4.48 in Nep3⁺; padj = 0.0011 and 3.8 × 10⁻¹⁶), TCF15 (log₂FC = 1.69 and 4.47; padj = 0.011 and 1.6 × 10⁻²¹), and GLI4 (log₂FC = 1.77 and 4.31; padj = 1.6 × 10⁻⁶ and 3.5 × 10⁻⁵⁴). Together, these results reveal both known and novel transcription factors with cell type restricted promoter activity, underscoring their potential roles in cnidocyte cell type specification **(Figures 5 and S5).**

**Figure 5.**
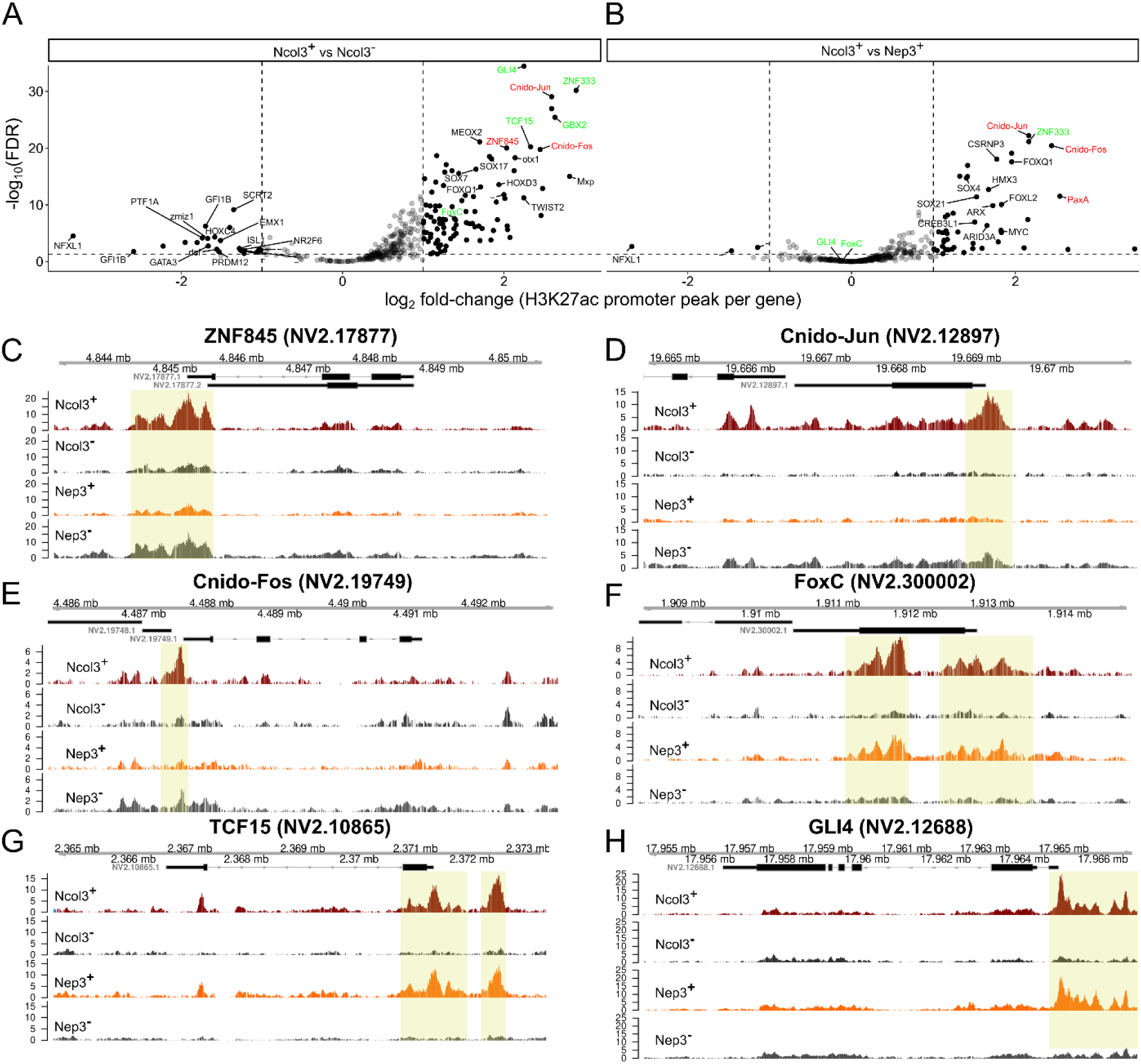
Promoter-centered H3K27ac profiling highlights cell type-specific transcription factor regulation in cnidocytes. **(A-B)** Volcano plots of differential H3K27ac signal at promoter-proximal peaks of TFs. Selected known TFs are shown in red and putative cnidocyte specific TFs are shown in green. **(A)** Comparison between Ncol3⁺ and Ncol3⁻ populations highlights transcription factors with increased acetylation in Ncol3⁺ cells, including ZNF845, Cnido-Fos, TCF15, and GLI4. **(B)** Comparison between Ncol3⁺ and Nep3⁺ cells reveals distinct regulatory programs, with enrichment of Cnido-Jun and FoxC acetylation in Ncol3⁺ cells. Each dot represents a promoter peak, with selected transcription factors labeled. **(C–H)** Genome browser tracks showing representative examples of transcription factor loci with cell type-specific promoter acetylation.

### Transgenic reporter assays validate regulatory elements

To functionally assess the activity of candidate regulatory elements identified by CUT&Tag, we generated transgenic reporter constructs and tested their regulatory potential *in vivo*. Approximately 1 kb sequences enriched in cnidocytes were cloned upstream of the mTurquoise2 fluorescent reporter and flanked by I-SceI meganuclease sites **(Figures 6A and S6)**. These 4 constructs, derived from regulatory regions near NV2.18817, NV2.13059, NV2.5200, NV2.5232 **(Figures S6A-D)**, were injected into *Ncol3::mOrange2* zygotes, and reporter activity was qualitatively assessed by fluorescence microscopy. Consistent with the CUT&Tag enrichment and cell type biased RNA expression of the associated genes **(Figure S6)**, all four elements drove reporter expression specifically in tentacle cnidocytes **(Figures 6B, 6C and S6E)**.

**Figure 6.**
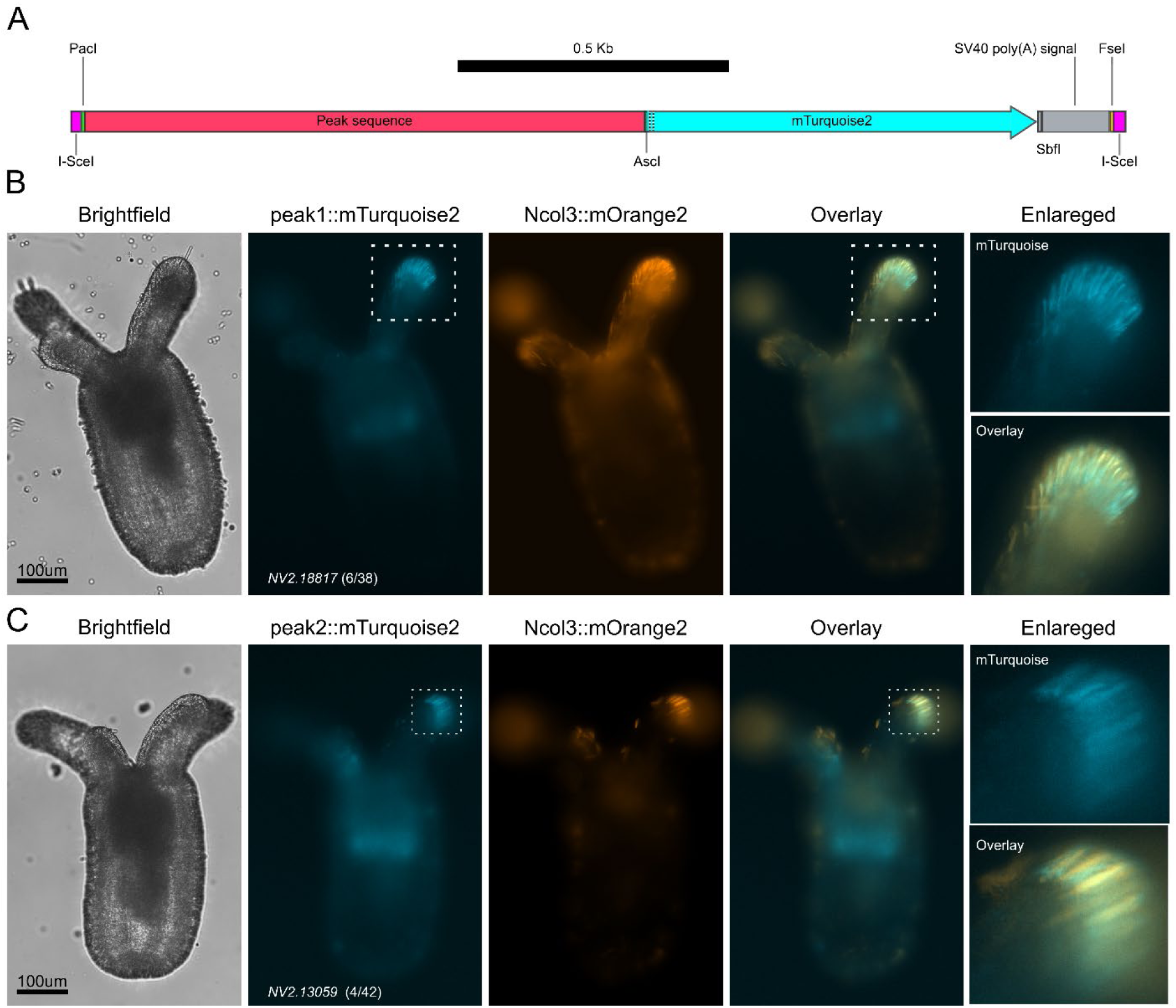
Functional validation of candidate regulatory elements in vivo. **(A)** Schematic of reporter construct design. ∼1 kb genomic sequences enriched in cnidocytes were cloned upstream of the mTurquoise2 fluorescent reporter and flanked by I-SceI meganuclease sites. **(B–C)** Reporter constructs were injected into *Ncol3::mOrange* zygotes, and fluorescence microscopy was used to assess activity. Scale bar, 100 µm. The area surrounding the tentacle tips (white rectangle) is enlarged on the right. Adjacent gene model and the number of positive animals (e.g., 6/38) are provided.

## Discussion

In this study, we provide the first cell type specific profiling of a histone modification in cnidocytes, the specialized stinging cells unique to cnidarians. By integrating CUT&Tag, RNA-seq, and transgenesis, we reveal the cis-regulatory signature that underlies cnidocyte cell type diversification. These results highlight how chromatin landscapes capture regulatory information that extends beyond transcriptomics and establish a framework for enhancer discovery in non-model organisms, paving the way for comparative epigenomics studies in early animals ^39^.

Our CUT&Tag profiling of H3K27ac revealed that Ncol3⁺ and Nep3⁺ cells are characterized by distinct enhancer modules **(Figures 4 and S4)**, separating shared regulatory programs from cell type restricted activity. Hierarchical clustering of acetylation profiles grouped putative enhancers into distinct modules **(Figure S4D)**, with certain clusters enriched specifically in nematocytes or spirocytes. These findings demonstrate that histone acetylation profiles can resolve subtle differences in cell identity, underscoring the importance of chromatin-level regulation in cnidocyte differentiation.

The analysis recovered several previously described transcription factors, including Cnido-Jun, Cnido-Fos, and ZNF845, validating our approach **(Figure 5)**. In addition, we identified zinc finger and homeobox factors such as ZNF333, GBX2, SCRT2, and PTF1A, which displayed highly restricted promoter activity **(Figure S5)**. Most strikingly, we uncovered a set of previously undescribed candidates that exhibited nematocyte-specific acetylation, including homologs of FoxC, TCF15, and GLI4 **(Figures 5F-H)**. The functional significance of these factors is supported by RNA-seq data, which showed strong upregulation in both Ncol3⁺ and Nep3⁺ cells (FoxC log₂FC = 3.06/4.48; TCF15 = 1.69/4.47; GLI4 = 1.77/4.31). The involvement of Fox, TCF, and Gli family members, key regulators of cell fate specification across Metazoa ^40-42^, suggests that cnidocyte differentiation is regulated by a combination of conserved transcription factors and lineage-specific regulators, reinforcing the view that new cell types emerge through usage of ancient regulatory modules ^43^.

Beyond transcription factors, our analysis highlighted candidate TF binding sites strongly enriched for H3K27ac **(Figure 4C)**. Four regions, located at proximal promoters, stood out due to their sharp acetylation peaks, enrichment of motifs such as HAP3, ROX8, and Jun-Fos, and the biased expression of their neighboring genes **(Figure S6)**. To directly test enhancer activity, we cloned ∼1 kb fragments upstream of an mTurquoise2 reporter and introduced them into *Ncol3::mOrange* zygotes. All constructs drove reporter expression specifically in tentacle cnidocytes, matching the CUT&Tag predictions **(Figures 6B, 6C and S6E)**. This validation demonstrates that cis-regulatory elements identified by chromatin profiling are capable of driving cell type specific transcription *in vivo*, providing a proof of principle for promoter/enhancer discovery and functional testing in *Nematostella vectensis*.

These findings also have broad evolutionary implications. Cnidocytes represent one of the most striking cellular innovations in animal evolution, and our results suggest that enhancer diversification is associated with the specification of cnidocyte types. This resembles the role of regulatory landscapes in controlling cell differentiation in neurons and blood cells in bilaterians ^44,45^. One of the most exciting implications of our findings is the potential they open for comparative epigenomics to shed light on how cell type regulatory programs evolve. We can now apply the same approach to compare chromatin landscapes across other cnidarians, early-diverging animals, and bilaterians, to assess which enhancer-TF modules are deeply conserved vs. those that are innovations. Recent work has shown, for example, that enhancers from evolutionarily distant species, as ancient as sponges, can share key motif grammars and drive similar developmental expression patterns in vertebrates ^46^. Likewise, our findings suggest conservation of the palindromic Jun-Fos motif and its corresponding TFs **(Figures 4C,5D and 5E).** Studies of epigenetic regulation in *Hydra* emphasize both conserved chromatin components and divergent regulatory strategies among cnidarians ^47^.

Methodologically, this work establishes a workflow combining epigenomic profiling, transcriptomics, and functional validation in an emerging cnidarian model system. The use of transgenic reporter lines provided the necessary cell type resolution to dissect regulatory regions in cnidocytes, representing a technical advance for functional genomics in early-branching animals. Looking forward, future studies should expand enhancer validation to a broader set of regulatory elements and employ quantitative approaches to assess their activity. High-throughput reporter assays and perturbation of candidate transcription factors will provide deeper mechanistic insight. Comparative studies across cnidarians and bilaterians could test whether the enhancer-TF combinations identified here reflect deeply conserved regulatory logic or lineage-specific innovations. Although our data supports nematocyte heterogeneity **(Figures 2F and G)**, single cell multi-omics approaches integrating chromatin accessibility, histone modifications, and transcriptional states within the same cells would further refine the connections between chromatin landscapes and transcriptional programs and elucidate cnidocyte diversity. Notably, recent single-cell sequencing studies in *Nematostella* have uncovered previously unrecognized cell types, including immune cells ^12,48^.

In summary, our study provides the first functional map of cnidocyte regulatory elements, linking chromatin landscapes, transcription factor activity, and enhancer function. By identifying both known and novel transcription factors and validating cis-regulatory elements *in vivo*, we provide a basis for understanding the chromatin mechanisms underlying cellular innovation in early-branching animals. These results underscore the central role of noncoding cis-regulatory DNA elements in driving cell type specialization and highlight cnidocytes as a powerful model for studying the evolution of regulatory mechanisms that generate novel cell types.

## Methods

### Animal culture and spawning

N. *vectensis* polyps were maintained at 18 °C in 16 ‰ sea salt water (Red Sea, Israel), used as the growth medium. Polyps were fed with Artemia salina nauplii three times a week. Spawning of gametes and fertilization were performed according to a published protocol ^49^ as follows: in brief, temperature was raised to 25 °C for 9 h and the animals were exposed to strong white light. Three hours after the induction, oocytes were mixed with sperm to allow fertilization. For cell sorting and imaging flow cytometry experiments, we used homozygous adult animals from the *Nep3::mOrange2* transgenic line ^50^ and heterozygous adults from the *Ncol3::mOrange2* line ^14^. Transgenic animals were visualized under an SMZ18 stereomicroscope equipped with a DS-Qi2 camera (Nikon, Japan) and Nikon TS100 equipped with a DS-Qi1Mc camera (Nikon).

### Cloning

Putative DNA regulatory regions of about 1Kb were identified by inspecting Ncol3^+^ and/or Nep3+ specific peaks using the Integrative Genomics Viewer (IGV, v2.16.2) ^51^. Only enriched motifs containing peaks with log_2_ fold change > 2 adjacent to a gene that also had a log_2_ fold change > 2 were considered. The selected regions flanked by PacI (upstream) and AscI (downstream) restriction sites were commercially synthesized and inserted into a vector pTwist Amp (Twist Bioscience, USA). This plasmid was then digested with PacI and AscI (New England Biolabs, USA) according to the manufacturer’s instructions. Digested fragments were size selected using 1% agarose gel and purified using Gel and PCR Clean-up kit (Macherey Nagel, Germany). Purified inserts were ligated into pCR2.1 vector, that was digested using the same restriction enzymes, using T4 DNA ligase (New England Biolabs), transformed into DH5α competent cells (New England Biolabs), and grown overnight in LB medium. For microinjections, plasmids were purified using Plasmid Plus Midi Kit (Qiagen, USA).

### Transgenic reporter lines

To validate regulatory elements, sperm release was induced in *Ncol3::mOrange* homozygous males, followed by fertilization of wild-type eggs. The resulting zygotes were microinjected with constructs using meganuclease-assisted delivery as previously described ^52^. 50ng ng/ml of plasmid DNA were used for each of the constructs. F_0_ 8 days old primary polyps were immobilized using 1/10 volume of 1 M MgCl₂ and scanned using a Nikon *Eclipse T100* inverted research microscope (Nikon) equipped with a Digital Sight *DS-Qi1Mc* camera and DS-U3 controller, using filter sets suitable for CFP and mOrange detection. Images were acquired with NIS-Elements BR, version 4.40.00 (Build 1084).

### Cell dissociation

Adult tentacles were enzymatically dissociated into single cells following established protocols with minor modifications ^53,54^. Briefly, adult polyps were relaxed for 30 minutes and then immobilized by adding 1/10 volume of 1 M MgCl₂ to the growth medium for 20 minutes. For each biological replicate, 5-6 individuals were used, with 4-5 replicates prepared per condition. Tentacles were dissected and collected into two 2-ml Eppendorf tubes, washed twice with one-third strength calcium/magnesium-free and EDTA-free artificial seawater (ASW; 3× stock: 17 mM Tris-HCl, 165 mM NaCl, 3.3 mM KCl, 9 mM NaHCO₃; final pH 8.0) to remove residual magnesium, and then incubated with 167 µg/ml LiberaseTM (Roche, Switzerland) in ASW at 37 °C for 30 minutes in a thermomixer (Eppendorf, Germany, 750 rpm) with occasional gentle pipetting until fully dissociated. The suspension was filtered through a 40-µm cell strainer (Corning, USA), centrifuged at 500 × g for 10 minutes at 4 °C, and resuspended in L-15-based culture medium supplemented with 2% heat-inactivated fetal bovine serum (FBS), 20 mM HEPES, and calcium/magnesium-free 10× PBS adjusted to 1.42× PBS concentration. Cell viability was assessed by adding Calcein Violet-AM (BioLegend, USA, 1:1000) and Zombie NIR (BioLegend, 1:250).

### Fluorescence-Activated Cell Sorting (FACS) of cell populations

FACSAria III (BD Biosciences, USA) equipped with 405 nm, 408 nm, 561 nm and 633 nm lasers was used to quantitively assess mOrange expression and to isolate cell populations based on mOrange expression status. Data was acquired using FACSDiva 9.0 (BD Biosciences). Per run, 30,000 events were recorded. WT animals were used to define mOrange positivity, and only Calcein Violet-AM positive, Zombie NIR negative viable cells were collected. To avoid RNase contamination, the FACS machine was sterilized by running bleach for 5 minutes followed by 5 minutes of 70 % ethanol at maximal speed. 100 µm nozzle was used for sorting. Immediately before sorting, cells were filtered using 5 mL round-bottom tubes with 35 µm cell strainer cap (Corning) and mixed vigorously by pipetting. For each run, 1-5 x 10^6^ cells/mL were used. Cells were sorted at low pressure at a speed of up to 3,000 events/sec. The sample tubes were kept at 4 °C throughout the process. To maximize yield by reducing cell adherence to the sides of the tubes, 15 mL conical bottom tubes were filled with PBS with 20 % fetal bovine serum (FBS) (Gibco) and incubated overnight at 4 °C degrees. Immediately before sorting, the tubes were emptied and filled with 1x cold PBS (calcium/magnesium free) (Hylabs). Collection tubes were kept at 4 °C during the sorting process. 600,000 viable cells were collected per cell fraction. Once completed, cells were centrifuged at 1000 × *g* at 4 °C for 10 minutes and immediately used for RNA extraction or for CUT&Tag. FCS files were further analyzed and visualized using FlowJo v10 (BD Biosciences) or FCS Express v7 (De Novo Software). Each measurement consisted of 3 or more biological replicates.

### RNA extraction

RNA was extracted from pelleted FACS-sorted cells using TRIzol Reagent (Thermo Fisher Scientific, USA) according to the manufacturer’s instructions with slight modifications. Briefly, cells were lysed in TRIzol by vigorous pipetting. During the precipitation step, an equal volume of isopropanol was added to the aqueous phase, together with 2 µL glycogen (Thermo Fisher Scientific) per sample. The mixture was inverted 10 times, incubated at -20 °C for 30 minutes, and centrifuged at maximum speed (∼21,000 × *g*) at 4 °C for 30 minutes. The resulting pellet was washed twice with 75% ethanol prepared in DEPC-treated water and resuspended in 15 µL nuclease-free water. RNA concentration was determined using the Qubit RNA High Sensitivity (HS) Assay Kit (Thermo Fisher Scientific), and RNA integrity was assessed with the Bioanalyzer RNA 6000 Pico Kit (Agilent, USA).

### RNA-seq library construction and sequencing

Only samples with an RNA integrity number (RIN) >7.5 were processed. Libraries were prepared from 100 ng of total RNA isolated from mOrange^+^ and mOrange^-^ cells of the *Nep3::mOrange* and *Ncol3::mOrange* transgenic lines. RNA-seq libraries were generated using the NEBNext Ultra II Directional RNA Library Prep Kit with Sample Purification Beads (New England Biolabs) according to the manufacturer’s protocol. Libraries were PCR-amplified and indexed using the NEBNext Multiplex Oligos for Illumina (New England Biolabs) with 14 amplification cycles. Amplified products were assessed with the High Sensitivity D1000 DNA Screen Tape Assay on an Agilent 2200 TapeStation system, and samples displaying a narrow distribution with a peak size of ∼300 bp were sequenced on a NextSeq 2000 platform (Illumina, USA) in dual-index mode with 120 bp single-end reads.

### CUT&Tag sample preparation

CUT&Tag was performed following the V3 protocol described by Kaya-Okur *et al.* ^37^ (Bench-top CUT&Tag V3; https://dx.doi.org/10.17504/protocols.io.bcuhiwt6). Briefly, for nuclei preparation, cells (6 × 10^5^ per reaction) were resuspended in freshly prepared NE1 buffer consisting of 20 mM HEPES-KOH (pH 7.9), 50 mM KCl, 5 mM spermidine, and 0.05% Triton X-100. After a 10 min incubation on ice, nuclei were pelleted by centrifugation (2000 × *g*, 5 min, 4 °C), the supernatant was removed, and nuclei were resuspended in wash buffer (20 mM HEPES pH 7.5, 150 mM NaCl, 0.5 mM spermidine, protease inhibitors). Nuclei were bound to Concanavalin A-coated magnetic beads (Cell Signaling Technology, cat #93569) and incubated (overnight at 4 °C) with anti-histone H3 (acetyl K27) antibody (Abcam, ChIP Grade cat. ab4729) or anti-IgG rabbit control (Sigma) followed by a secondary goat anti-rabbit unconjugated antibody (Jackson ImmunoResearch, USA) for 1 h at room temperature. Bead-bound nuclei were then incubated with pA-Tn5 transposase (EpyCypher, USA) for 1 h at room temperature. After stringent washes, tagmentation was initiated by addition of MgCl₂ (10 mM final) for 1 h at 37 °C. DNA was extracted using phenol-chloroform, PCR-amplified using 14 cycles, and libraries were purified using Mag-Bind TotalPure NGS beads (Omega Biotek, USA) prior to sequencing. Amplified products were assessed with the High Sensitivity D1000 DNA Screen Tape Assay on an Agilent 2200 TapeStation system, and samples displaying mono and di-nucleosome enrichment with a strong peak size of ∼300 bp were sequenced on a NextSeq 2000 platform in dual-index mode with R1 80bp and R2 40bp paired-end reads.

### Imaging flow cytometry data acquisition

Dissociated tentacle-derived cells were analyzed using an Amnis ImageStreamX Mk II imaging cytometer (Luminex, USA) equipped with 405, 488, 561, and 642 nm lasers, 12 acquisition channels, and two cameras, with a 60× objective operated in low-flow/high-sensitivity mode. Data were acquired with INSPIRE software (Luminex) using the following configuration: Ch01 (bright field, camera 1), Ch03 (mOrange), Ch06 (side scatter, SSC), Ch07 (Calcein Violet-AM), Ch09 (bright field, camera 2), and Ch12 (Zombie NIR). Cells were resuspended in 100 μL PBS in 1.5 mL tubes, and approximately 1 × 10⁷ cells were loaded per sample. For each sample, 10,000 focused singlet events were recorded. Focused events were identified using the Gradient RMS parameter in Ch01 and Ch09, while singlets were gated based on event area (X-axis) against aspect ratio (Y-axis) in Ch01, retaining low-aspect-ratio events to capture mature cnidocytes. Downstream analyses were restricted to this population. Image analysis was performed using IDEAS 6.3 software (Luminex). Circularity was quantified with the Shape Change Wizard, which computes a circularity score for each population by measuring deviations of the mask from a perfect circle.

### Statistical analysis, clustering, and dimensionality reduction of Imaging cytometry data

Imaging flow cytometry data from two conditions (Ncol3⁺ and Nep3⁺) were imported into Rstudio (v4.4.1). Non-numeric metadata columns and mOrange-related fluorescence features (Ch03 channel) were excluded to avoid bias from reporter signal. Zero-variance features were removed, and the remaining features (n=545) were z-score scaled prior to downstream analysis.

Principal component analysis (PCA) was performed using the stats4 package, and the first two principal components were visualized. Unsupervised clustering was carried out with k-means clustering (*cluster* v2.1.6 package), with the number of clusters (k = 4) determined empirically. Cluster assignments were overlaid on PCA plots, with representative cells from each cluster labeled for reference.

To capture non-linear structure in the data, uniform manifold approximation and projection (UMAP) was performed using the uwot v0.2.2 package (parameters: *n_neighbors* = 15, *min_dist* = 0.1, Euclidean metric). K-means clustering (k = 4) was applied to the scaled feature matrix, and the resulting cluster assignments were visualized on UMAP coordinates. Selected object numbers were highlighted to aid interpretation. Cluster composition was quantified by merging cluster assignments with sample metadata. Proportional analysis was performed by computing the distribution of clusters within each condition. These proportions were displayed as stacked bar plots generated with ggplot2 v3.5.1.

For selected morphological features (e.g., elongatedness, circularity, aspect ratio etc.), pairwise comparisons between conditions were performed using two-sample *t*-tests (ggpubr v0.6.0, function stat_compare_means, Welch correction). Results were displayed on violin plots overlaid with internal boxplots and jittered points (ggplot2), with medians indicated by a red marker (stat_summary(fun = median)). Significance codes were annotated directly on the plots to facilitate interpretation.

### Bulk RNA-seq analysis and GO functional analysis

The quality of raw reads was assessed and visualized using FastQC v0.12.0 ^55^, with summary reports generated through multiqc v1.26 ^56^. Reads were trimmed using Trimmomatic v0.39 ^57^ with the following parameters: ILLUMINACLIP:TruSeq3-SE:2:30:10, LEADING:3, TRAILING:3, SLIDINGWINDOW:4:15, and MINLEN:36. The trimmed reads were aligned to the Nv2 *N. vectensis* reference genome ^58^, supplemented with the mOrange transgene sequence, using STAR v2.7.10a ^59^. Gene-level read counts were summarized using featureCounts v2.0.1 ^60^. Genes with read counts >= 50 across all samples were kept. Differentially expressed genes (DEGs) across different cell fractions were identified using DESeq2 v1.44.0 ^61^ with the design = ∼ condition argument. DEGs were defined as genes with an absolute log2 fold change > 1 and an adjusted p-value < 0.05. Principal component analysis (PCA) was performed on variance-stabilized transformed (vst) data using the DESeq2 vst function with the blind = FALSE argument.

Gene Ontology (GO) enrichment comparison across the three sets (shared, correlated, anti-correlated) was performed with clusterProfiler v4.12.6 (function compareCluster, method = "enricher") using a custom molecular function and cellular function annotations. Parameters were pvalueCutoff = 0.05, pAdjustMethod = "BH", qvalueCutoff = 0.20, and gene-set size bounds minGSSize = 10, maxGSSize = 500. The resulting compareCluster object was converted to a data frame for export and visualization. To present human-readable terms, GO IDs were mapped to names with GO.db v3.19.1 and AnnotationDbi v1.66.0. Missing/obsolete IDs were returned as NAs. The Description field in the compareCluster result was replaced by these names and entries with missing names were dropped prior to plotting. Visualization used ggplot2 v3.5.1 and viridis v0.6.5. Enrichment comparisons were displayed as dot plots, with point size proportional to gene count and color encoding the significance level as -log10(p.adjust).

### Identification of transcriptomic gene modules by unsupervised clustering

Raw read count matrices were imported into Rstudio and processed using the tidyverse v2.0.0. Non-informative columns (chromosome, start, end, strand) and one low-quality replicate (Ncol3_neg_4) were removed. Genes with fewer than 50 total counts across all samples were filtered out. Differential expression analysis was performed with DESeq2 (design= ∼condition), and a likelihood ratio test (LRT) was used to identify genes whose expression varied significantly across conditions (padj < 0.05). Variance-stabilizing transformation was applied to normalized counts, and for each gene, average expression per condition was calculated. Expression values were centered and scaled by gene. Fuzzy c-means clustering was performed with the e1071 v1.7-16 package (centers = 4, fuzziness parameter *m* = 2), grouping genes into four clusters. Genes were ordered by cluster assignment, and scaled expression values were visualized as a heatmap with the superheat v1.0.0 R package ^62^. A continuous *cividis* palette with 500 color levels (viridis::cividis(500)) was applied, with columns hierarchically clustered and rows fixed by cluster order. The resulting heatmap displays all genes significantly varying across conditions (LRT padj < 0.05).

### CUT&Tag raw data processing, alignment and downstream analysis

Quality assessment and visualization of the CUT&Tag raw reads were performed using FastQC v0.12.0 ^55^, and summary reports were compiled with MultiQC v1.26 ^56^. Adapter and quality trimming of paired-end CUT&Tag reads was performed using Cutadapt (v5.0, Python 3.12.9) ^63^. Sequencing adapters were removed from both R1 and R2 reads using the following sequences: Tn5ME-A: 5ʹ-TCGTCGGCAGCGTCAGATGTGTATAAGAGACAG-3ʹ, Tn5ME-B: 5ʹ-GTCTCGTGGGCTCGGAGATGTGTATAAGAGACAG-3ʹ, Tn5ME-rev: 5ʹ-CTGTCTCTTATACACATCT-3ʹ. Bases with Phred quality scores below 20 and terminal ambiguous nucleotides (N) were trimmed. Read pairs were discarded if either pair failed filtering (--pair-filter=any), and reads shorter than 20 bp after trimming were discarded. Trimmed paired-end reads were then aligned to the Nv2 *N. vectensis* reference genome ^58^ using Bowtie2 (v2.5.2). This resulted in overall alignment rate > 95 % for all samples and sequencing depth ranging from 3.2 to 16.2 million reads per sample.

Mapped fragment size distributions were assessed to verify CUT&Tag library quality. This confirmed nucleosomal patterns and the expected nucleosomal length periodicity (about 150bp). Properly paired reads were filtered using a minimal quality score (-q) =2 and extracted using *samtools* v1.19.2 ^64^. Peaks were called using MACS2 v 2.2.9.1 ^65^ (paired-end mode) with a q-value threshold of 0.1 to control for false discovery. Normalized coverage tracks (bigWig files) were produced from aligned BAM files using *deepTools* v3.5.5 ^66^ with RPGC normalization (reads per genomic content), enabling accurate comparisons across samples. These tracks were used for genome browser inspection and visualization using IGV v2.16.2 ^51^ and Gviz v1.52.0 ^67^ as well as for computing signal matrices around peak summits. Heatmaps and average profile plots were subsequently generated with the *deepTools* functions computeMatrix, plotHeatmap, plotProfile to visualize genome-wide signal distributions.

To quantify CUT&Tag signal across samples, a count matrix was generated. First, a unified peak set was constructed by merging condition-specific peaks across all samples using bedtools v2.31.1 ^68^ *merge*, producing a non-redundant consensus peak list. Read counts per peak were then obtained using featureCounts v2.0.1, with parameters set to count properly paired reads overlapping the merged peak coordinates. This yielded a peak-by-sample count matrix, which served as the input for downstream statistical analyses.

Differential enrichment analysis was performed using DESeq2 ^61^, with size factor normalization applied to account for differences in sequencing depth. Variance-stabilizing transformation (VST) was used to generate normalized expression values for exploratory analyses including principal component analysis (PCA) and hierarchical clustering, which revealed clear sample grouping by cell type. Significantly differential peaks (padj < 0.05) were further visualized as clustered heatmaps using *pheatmap* v1.0.12 ^69^ and *ComplexHeatmap* v2.20.0 ^70^, highlighting cell type-specific modules. Motif enrichment analysis was performed using the HOMER suite v5.1^71^. Input sequences were provided as FASTA files of enriched regions. For de novo motif discovery, we ran findMotifs.pl on the FASTA inputs, specifying multiple motif lengths (8, 10, and 12 bp). Analyses were performed with default parameters.

Finally, peaks were annotated relative to genomic features using *ChIPseeker* v1.40.0 ^72^, and peak-associated genes were subjected to gene ontology enrichment analysis using *ClusterProfiler* to gain functional insights into the regulatory programs marked by H3K27ac.

### scRNA-seq expression analysis

We complemented our transcriptomic profiling with a recently published single-cell RNA-seq atlas ^12^. The cnidocyte subset .Robj file was obtained from the publicly available UCSC Cell Browser “Sea Anemone Atlas” (https://cells.ucsc.edu/?ds=sea-anemone-atlas) and loaded into Rstudio. To highlight putative spirocyte markers, we visualized the top Ncol3⁺ upregulated genes that were downregulated in Nep⁺ cells using the DotPlot function in Seurat v4.1.1 ^73^.

## Data availability

Raw sequencing data have been deposited in the NCBI Sequence Read Archive (SRA) under BioProject accessions PRJNA1333423 (CUT&Tag) and PRJNA1333351 (RNA-seq). Processed data, including normalized signal tracks (bigWig), MACS2 narrowPeak files, and the RNA-seq gene count matrix, have been deposited in the Gene Expression Omnibus (GEO) under accessions GSE309459 (CUT&Tag) and GSE309458 (RNA-seq).

## Acknowledgements

We are grateful to Ms. Adi Turjeman and Dr. Michal Bronstein of the Center for Genomic Technologies, The Hebrew University of Jerusalem, for their help with sequencing. We are also grateful to Dr. Hadas Segev-Yekutiel of the Core Research Facility, Faculty of Medicine, The Hebrew University of Jerusalem, Ein Kerem, for support with imaging flow cytometry data acquisition, and to Dr. Ola Karmi of the Research Infrastructure Facility, The Hebrew University of Jerusalem, for her assistance with imaging flow cytometry data analysis and FACS. We thank Dr. Reuven Aharoni (The Hebrew University of Jerusalem) for his technical assistance. This work was supported by a European Research Council Consolidator Grant (AntiViralEvo; 863809) and as well as grant from the Israel Science Foundation (636/21) to YM.

## Notes

### Competing Interest Statement

The authors have declared no competing interest.

## References

1. Babonis, L.S. (2025). That stinging sensation: modularity and the origin of the stinging cell. Integrative And Comparative Biology, icaf070.

2. Fautin, D.G. (2009). Structural diversity, systematics, and evolution of cnidae. Toxicon 54, 1054–1064. 10.1016/j.toxicon.2009.02.024.

3. Beckmann, A., and Ozbek, S. (2012). The nematocyst: a molecular map of the cnidarian stinging organelle. Int J Dev Biol 56, 577–582. 10.1387/ijdb.113472ab.

4. Fautin, D.G. (2009). Structural diversity, systematics, and evolution of cnidae. Toxicon 54, 1054–1064.

5. Kass-Simon, G., and Scappaticci, J., AA (2002). The behavioral and developmental physiology of nematocysts. Canadian Journal of Zoology 80, 1772–1794.

6. Babonis, L.S., Enjolras, C., Reft, A.J., Foster, B.M., Hugosson, F., Ryan, J.F., Daly, M., and Martindale, M.Q. (2023). Single-cell atavism reveals an ancient mechanism of cell type diversification in a sea anemone. Nature Communications 14, 885.

7. Siebert, S., Farrell, J.A., Cazet, J.F., Abeykoon, Y., Primack, A.S., Schnitzler, C.E., and Juliano, C.E. (2019). Stem cell differentiation trajectories in Hydra resolved at single-cell resolution. Science 365, eaav9314.

8. Holstein, T., and Tardent, P. (1984). An ultrahigh-speed analysis of exocytosis: nematocyst discharge. Science 223, 830–833.

9. Babonis, L.S., and Martindale, M.Q. (2014). Old cell, new trick? Cnidocytes as a model for the evolution of novelty. American Zoologist 54, 714–722.

10. Karabulut, A., McClain, M., Rubinstein, B., Sabin, K.Z., McKinney, S.A., and Gibson, M.C. (2022). The architecture and operating mechanism of a cnidarian stinging organelle. Nature Communications 13, 3494.

11. Özbek, S., Balasubramanian, P.G., and Holstein, T.W. (2009). Cnidocyst structure and the biomechanics of discharge. Toxicon 54, 1038–1045.

12. Cole, A.G., Steger, J., Hagauer, J., Denner, A., Ferrer Murguia, P., Knabl, P., Narayanaswamy, S., Wick, B., Montenegro, J.D., and Technau, U. (2024). Updated single cell reference atlas for the starlet anemone Nematostella vectensis. Frontiers in Zoology 21, 8.

13. Sebé-Pedrós, A., Saudemont, B., Chomsky, E., Plessier, F., Mailhé, M.-P., Renno, J., Loe-Mie, Y., Lifshitz, A., Mukamel, Z., and Schmutz, S. (2018). Cnidarian cell type diversity and regulation revealed by whole-organism single-cell RNA-Seq. Cell 173, 1520–1534. e1520.

14. Sunagar, K., Columbus-Shenkar, Y.Y., Fridrich, A., Gutkovich, N., Aharoni, R., and Moran, Y. (2018). Cell type-specific expression profiling unravels the development and evolution of stinging cells in sea anemone. BMC biology 16, 108.

15. Babonis, L.S., and Martindale, M.Q. (2017). PaxA, but not PaxC, is required for cnidocyte development in the sea anemone Nematostella vectensis. Evodevo 8, 14.

16. Steger, J., Cole, A.G., Denner, A., Lebedeva, T., Genikhovich, G., Ries, A., Reischl, R., Taudes, E., Lassnig, M., and Technau, U. (2022). Single-cell transcriptomics identifies conserved regulators of neuroglandular lineages. Cell Reports 40.

17. Tournière, O., Dolan, D., Richards, G.S., Sunagar, K., Columbus-Shenkar, Y.Y., Moran, Y., and Rentzsch, F. (2020). NvPOU4/Brain3 functions as a terminal selector gene in the nervous system of the cnidarian Nematostella vectensis. Cell reports 30, 4473–4489. e4475.

18. Babonis, L.S., Enjolras, C., Ryan, J.F., and Martindale, M.Q. (2022). A novel regulatory gene promotes novel cell fate by suppressing ancestral fate in the sea anemone Nematostella vectensis. Proceedings of the National Academy of Sciences 119, e2113701119.

19. Fridrich, A., Salinas-Saaverda, M., Kozlolvski, I., Surm, J.M., Chrysostomou, E., Tripathi, A.M., Frank, U., and Moran, Y. (2023). An ancient pan-cnidarian microRNA regulates stinging capsule biogenesis in Nematostella vectensis. Cell Reports 42.

20. Klemm, S.L., Shipony, Z., and Greenleaf, W.J. (2019). Chromatin accessibility and the regulatory epigenome. Nature Reviews Genetics 20, 207–220.

21. Heintzman, N.D., Hon, G.C., Hawkins, R.D., Kheradpour, P., Stark, A., Harp, L.F., Ye, Z., Lee, L.K., Stuart, R.K., and Ching, C.W. (2009). Histone modifications at human enhancers reflect global cell-type-specific gene expression. Nature 459, 108–112.

22. Creyghton, M.P., Cheng, A.W., Welstead, G.G., Kooistra, T., Carey, B.W., Steine, E.J., Hanna, J., Lodato, M.A., Frampton, G.M., and Sharp, P.A. (2010). Histone H3K27ac separates active from poised enhancers and predicts developmental state. Proceedings of the National Academy of Sciences 107, 21931–21936.

23. Ma, S., Zhang, B., LaFave, L.M., Earl, A.S., Chiang, Z., Hu, Y., Ding, J., Brack, A., Kartha, V.K., and Tay, T. (2020). Chromatin potential identified by shared single-cell profiling of RNA and chromatin. Cell 183, 1103–1116. e1120.

24. Bartosovic, M., Kabbe, M., and Castelo-Branco, G. (2021). Single-cell CUT&Tag profiles histone modifications and transcription factors in complex tissues. Nature biotechnology 39, 825–835.

25. Li, Y.E., Preissl, S., Miller, M., Johnson, N.D., Wang, Z., Jiao, H., Zhu, C., Wang, Z., Xie, Y., and Poirion, O. (2023). A comparative atlas of single-cell chromatin accessibility in the human brain. Science 382, eadf7044.

26. Ranzoni, A.M., Tangherloni, A., Berest, I., Riva, S.G., Myers, B., Strzelecka, P.M., Xu, J., Panada, E., Mohorianu, I., and Zaugg, J.B. (2021). Integrative single-cell RNA-seq and ATAC-seq analysis of human developmental hematopoiesis. Cell stem cell 28, 472–487. e477.

27. Park, J., Lee, K., Kim, K., and Yi, S.-J. (2022). The role of histone modifications: from neurodevelopment to neurodiseases. Signal transduction and targeted therapy 7, 217.

28. Delás, M.J., Kalaitzis, C.M., Fawzi, T., Demuth, M., Zhang, I., Stuart, H.T., Costantini, E., Ivanovitch, K., Tanaka, E.M., and Briscoe, J. (2023). Developmental cell fate choice in neural tube progenitors employs two distinct cis-regulatory strategies. Developmental cell 58, 3–17. e18.

29. Schwaiger, M., Schönauer, A., Rendeiro, A.F., Pribitzer, C., Schauer, A., Gilles, A.F., Schinko, J.B., Renfer, E., Fredman, D., and Technau, U. (2014). Evolutionary conservation of the eumetazoan gene regulatory landscape. Genome research 24, 639–650.

30. Cazet, J.F., Siebert, S., Little, H.M., Bertemes, P., Primack, A.S., Ladurner, P., Achrainer, M., Fredriksen, M.T., Moreland, R.T., and Singh, S. (2023). A chromosome-scale epigenetic map of the Hydra genome reveals conserved regulators of cell state. Genome research 33, 283–298.

31. Al-Shaer, L., Havrilak, J., and Layden, M.J. (2021). Nematostella vectensis as a Model System. In Handbook of marine model organisms in experimental biology, (CRC Press), pp. 107–128.

32. Rentzsch, F., and Technau, U. (2016). Genomics and development of Nematostella vectensis and other anthozoans. Current opinion in genetics & development 39, 63–70.

33. Layden, M.J., Rentzsch, F., and Röttinger, E. (2016). The rise of the starlet sea anemone Nematostella vectensis as a model system to investigate development and regeneration. Wiley Interdisciplinary Reviews: Developmental Biology 5, 408–428.

34. Zenkert, C., Takahashi, T., Diesner, M.-O., and Özbek, S. (2011). Morphological and molecular analysis of the Nematostella vectensis cnidom. PloS one 6, e22725.

35. Columbus-Shenkar, Y.Y., Sachkova, M.Y., Macrander, J., Fridrich, A., Modepalli, V., Reitzel, A.M., Sunagar, K., and Moran, Y. (2018). Dynamics of venom composition across a complex life cycle. Elife 7, e35014.

36. Zuba-Surma, E.K., Kucia, M., Abdel-Latif, A., Lillard, J.W., and Ratajczak, M.Z. (2007). The ImageStream System: a key step to a new era in imaging. Folia histochemica et cytobiologica 45, 279–290.

37. Kaya-Okur, H.S., Wu, S.J., Codomo, C.A., Pledger, E.S., Bryson, T.D., Henikoff, J.G., Ahmad, K., and Henikoff, S. (2019). CUT&Tag for efficient epigenomic profiling of small samples and single cells. Nature communications 10, 1930.

38. Landt, S.G., Marinov, G.K., Kundaje, A., Kheradpour, P., Pauli, F., Batzoglou, S., Bernstein, B.E., Bickel, P., Brown, J.B., and Cayting, P. (2012). ChIP-seq guidelines and practices of the ENCODE and modENCODE consortia. Genome research 22, 1813–1831.

39. Xiao, S., Xie, D., Cao, X., Yu, P., Xing, X., Chen, C.-C., Musselman, M., Xie, M., West, F.D., and Lewin, H.A. (2012). Comparative epigenomic annotation of regulatory DNA. Cell 149, 1381–1392.

40. Yasuoka, Y., Matsumoto, M., Yagi, K., and Okazaki, Y. (2020). Evolutionary history of GLIS genes illuminates their roles in cell reprograming and ciliogenesis. Molecular biology and evolution 37, 100–109.

41. Shimeld, S.M., Boyle, M.J., Brunet, T., Luke, G.N., and Seaver, E.C. (2010). Clustered Fox genes in lophotrochozoans and the evolution of the bilaterian Fox gene cluster. Developmental biology 340, 234–248.

42. Rodriguez-Fraticelli, A.E., Weinreb, C., Wang, S.-W., Migueles, R.P., Jankovic, M., Usart, M., Klein, A.M., Lowell, S., and Camargo, F.D. (2020). Single-cell lineage tracing unveils a role for TCF15 in haematopoiesis. Nature 583, 585–589.

43. Arendt, D., Musser, J.M., Baker, C.V., Bergman, A., Cepko, C., Erwin, D.H., Pavlicev, M., Schlosser, G., Widder, S., and Laubichler, M.D. (2016). The origin and evolution of cell types. Nature Reviews Genetics 17, 744–757.

44. Lara-Astiaso, D., Weiner, A., Lorenzo-Vivas, E., Zaretsky, I., Jaitin, D.A., David, E., Keren-Shaul, H., Mildner, A., Winter, D., and Jung, S. (2014). Chromatin state dynamics during blood formation. science 345, 943–949.

45. Yao, B., Christian, K.M., He, C., Jin, P., Ming, G.-l., and Song, H. (2016). Epigenetic mechanisms in neurogenesis. Nature reviews neuroscience 17, 537–549.

46. Wong, E.S., Zheng, D., Tan, S.Z., Bower, N.I., Garside, V., Vanwalleghem, G., Gaiti, F., Scott, E., Hogan, B.M., and Kikuchi, K. (2020). Deep conservation of the enhancer regulatory code in animals. Science 370, eaax8137.

47. Pillai, A., Gungi, A., Reddy, P.C., and Galande, S. (2021). Epigenetic regulation in Hydra: Conserved and divergent roles. Frontiers in Cell and Developmental Biology 9, 663208.

48. Kozlovski, I., Sharoni, T., Levy, S., Jaimes-Becerra, A., Talice, S., Kwak, H.-J., Aleshkina, D., Grau-Bové, X., Karmi, O., and Rosental, B. (2025). Functional characterization of immune cells in a cnidarian reveals an ancestral antiviral program. bioRxiv, 2025.2001. 2024.634691.

49. Genikhovich, G., and Technau, U. (2009). Induction of spawning in the starlet sea anemone Nematostella vectensis, in vitro fertilization of gametes, and dejellying of zygotes. Cold Spring Harbor Protocols 2009, pdb. prot5281.

50. Columbus-Shenkar, Y.Y., Sachkova, M.Y., Macrander, J., Fridrich, A., Modepalli, V., Reitzel, A.M., Sunagar, K., and Moran, Y. (2018). Dynamics of venom composition across a complex life cycle. Elife 7. 10.7554/eLife.35014.

51. Thorvaldsdóttir, H., Robinson, J.T., and Mesirov, J.P. (2013). Integrative Genomics Viewer (IGV): high-performance genomics data visualization and exploration. Briefings in bioinformatics 14, 178–192.

52. Renfer, E., and Technau, U. (2017). Meganuclease-assisted generation of stable transgenics in the sea anemone Nematostella vectensis. Nature protocols 12, 1844–1854.

53. Admoni, Y., Kozlovski, I., Lewandowska, M., and Moran, Y. (2020). TATA Binding Protein (TBP) Promoter Drives Ubiquitous Expression of Marker Transgene in the Adult Sea Anemone Nematostella vectensis. Genes 11, 1081. 10.3390/genes11091081.

54. Kozlovski, I., Jaimes-Becerra, A., Sharoni, T., Lewandowska, M., Karmi, O., and Moran, Y. (2024). Induction of apoptosis by double-stranded RNA was present in the last common ancestor of cnidarian and bilaterian animals. PLoS Pathog 20, e1012320. 10.1371/journal.ppat.1012320.

55. Andrews, S. (2010). FastQC: A Quality Control Tool for High Throughput Sequence Data (Babraham Bioinformatics, Babraham Institute).

56. Ewels, P., Magnusson, M., Lundin, S., and Kaller, M. (2016). MultiQC: summarize analysis results for multiple tools and samples in a single report. Bioinformatics 32, 3047–3048. 10.1093/bioinformatics/btw354.

57. Bolger, A.M., Lohse, M., and Usadel, B. (2014). Trimmomatic: a flexible trimmer for Illumina sequence data. Bioinformatics 30, 2114–2120. 10.1093/bioinformatics/btu170.

58. Zimmermann, B., Montenegro, J.D., Robb, S.M., Fropf, W.J., Weilguny, L., He, S., Chen, S., Lovegrove-Walsh, J., Hill, E.M., and Chen, C.-Y. (2023). Topological structures and syntenic conservation in sea anemone genomes. Nature Communications 14, 8270.

59. Dobin, A., Davis, C.A., Schlesinger, F., Drenkow, J., Zaleski, C., Jha, S., Batut, P., Chaisson, M., and Gingeras, T.R. (2013). STAR: ultrafast universal RNA-seq aligner. Bioinformatics 29, 15–21. 10.1093/bioinformatics/bts635.

60. Liao, Y., Smyth, G.K., and Shi, W. (2014). featureCounts: an efficient general purpose program for assigning sequence reads to genomic features. Bioinformatics 30, 923–930. 10.1093/bioinformatics/btt656.

61. Love, M.I., Huber, W., and Anders, S. (2014). Moderated estimation of fold change and dispersion for RNA-seq data with DESeq2. Genome Biol 15, 550. 10.1186/s13059-014-0550-8.

62. Barter, R.L., and Yu, B. (2018). Superheat: An R package for creating beautiful and extendable heatmaps for visualizing complex data. Journal of Computational and Graphical Statistics 27, 910–922.

63. Martin, M. (2011). Cutadapt removes adapter sequences from high-throughput sequencing reads. EMBnet. journal 17, 10–12.

64. Li, H., Handsaker, B., Wysoker, A., Fennell, T., Ruan, J., Homer, N., Marth, G., Abecasis, G., Durbin, R., and Subgroup, G.P.D.P. (2009). The sequence alignment/map format and SAMtools. bioinformatics 25, 2078–2079.

65. Feng, J., Liu, T., Qin, B., Zhang, Y., and Liu, X.S. (2012). Identifying ChIP-seq enrichment using MACS. Nature protocols 7, 1728–1740.

66. Ramírez, F., Dündar, F., Diehl, S., Grüning, B.A., and Manke, T. (2014). deepTools: a flexible platform for exploring deep-sequencing data. Nucleic acids research 42, W187–W191.

67. Hahne, F., and Ivanek, R. (2016). Visualizing genomic data using Gviz and bioconductor. In Statistical genomics: methods and protocols, (Springer), pp. 335–351.

68. Quinlan, A.R., and Hall, I.M. (2010). BEDTools: a flexible suite of utilities for comparing genomic features. Bioinformatics 26, 841–842.

69. Kolde, R., and Kolde, M.R. (2015). Package ‘pheatmap’. R package 1, 790.

70. Gu, Z. (2022). Complex heatmap visualization. Imeta 1, e43.

71. Heinz, S., Benner, C., Spann, N., Bertolino, E., Lin, Y.C., Laslo, P., Cheng, J.X., Murre, C., Singh, H., and Glass, C.K. (2010). Simple combinations of lineage-determining transcription factors prime cis-regulatory elements required for macrophage and B cell identities. Molecular cell 38, 576–589.

72. Yu, G., Wang, L.-G., and He, Q.-Y. (2015). ChIPseeker: an R/Bioconductor package for ChIP peak annotation, comparison and visualization. Bioinformatics 31, 2382–2383.

73. Butler, A., Hoffman, P., Smibert, P., Papalexi, E., and Satija, R. (2018). Integrating single-cell transcriptomic data across different conditions, technologies, and species. Nature biotechnology 36, 411–420.

