## Supplementary Figures for "Epigenomic and transcriptomic analyses reveal cnidocyte specialization in a sea anemone"

### for

**Table of contents**

Supplementary Figure S1-S6

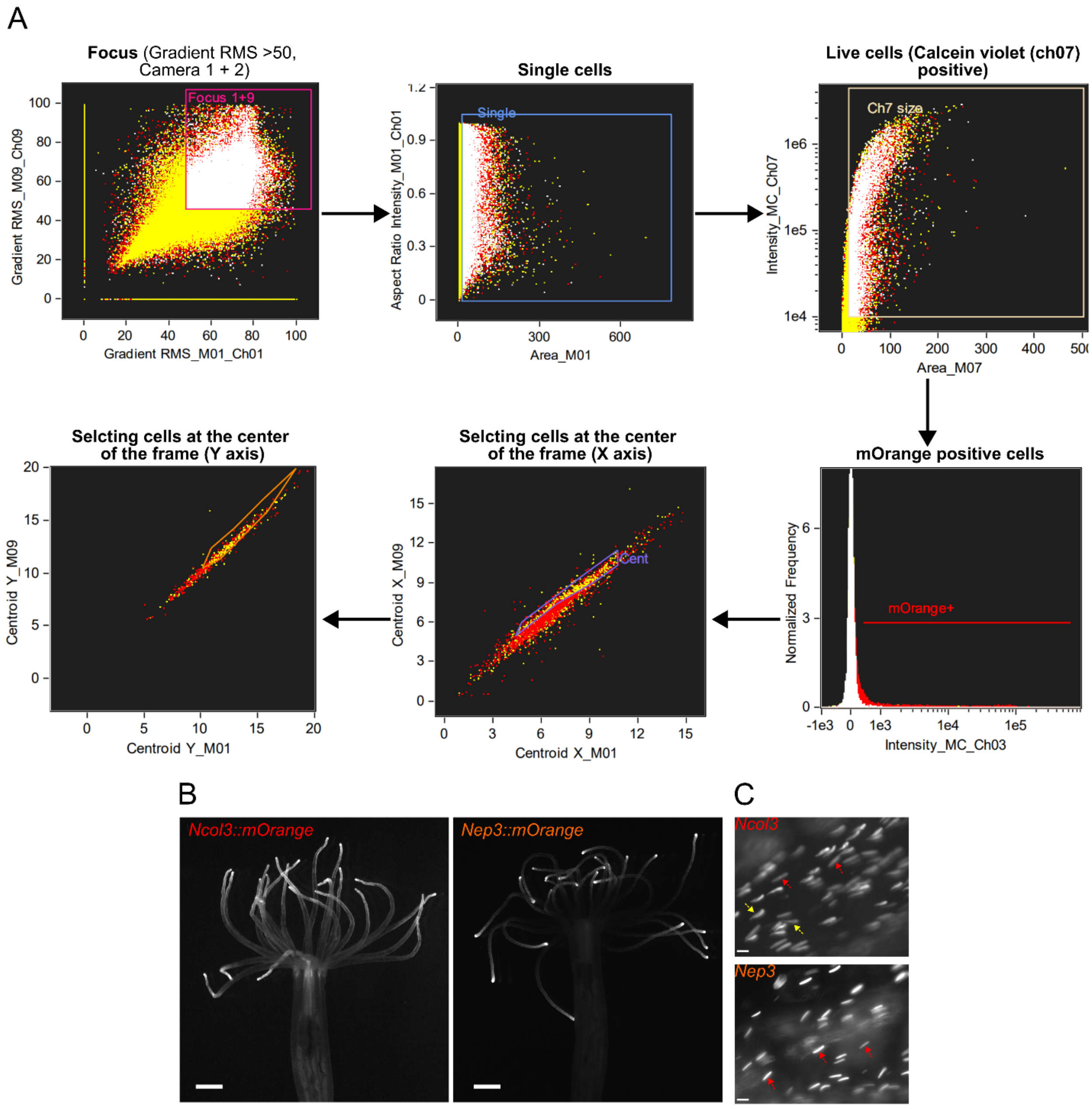


**Figure S1. Imaging and gating strategy for morphological analysis of cnidocytes. (A)** Gating strategy for imaging flow cytometry (ImageStream) used for morphological analysis of cnidocyte types. Sequential gates were applied to select focused cells (gradient RMS >50), single cells (aspect ratio vs. area), and live cells (Calcein violet positive, Ch07). From this population, mOrange$^{+}$ cells were identified, and off-center events were excluded by gating on centroid positions along the X and Y axes. **(B)** Representative fluorescence images of *Nematostella vectensis* polyps expressing the reporter constructs *Ncol3::mOrange* (left) and *Nep3::mOrange* (right). Scale bars: 1 mm. **(C)** Microscopy images of intact tentacles. In *Ncol3::mOrange* animals (top), both mOrange-positive nematocytes (red arrowheads) and spirocytes (yellow arrows) were observed. In contrast, *Nep3::mOrange* animals (bottom) contained mOrange-positive nematocytes only (red arrowheads). Scale bars: 10 µm.

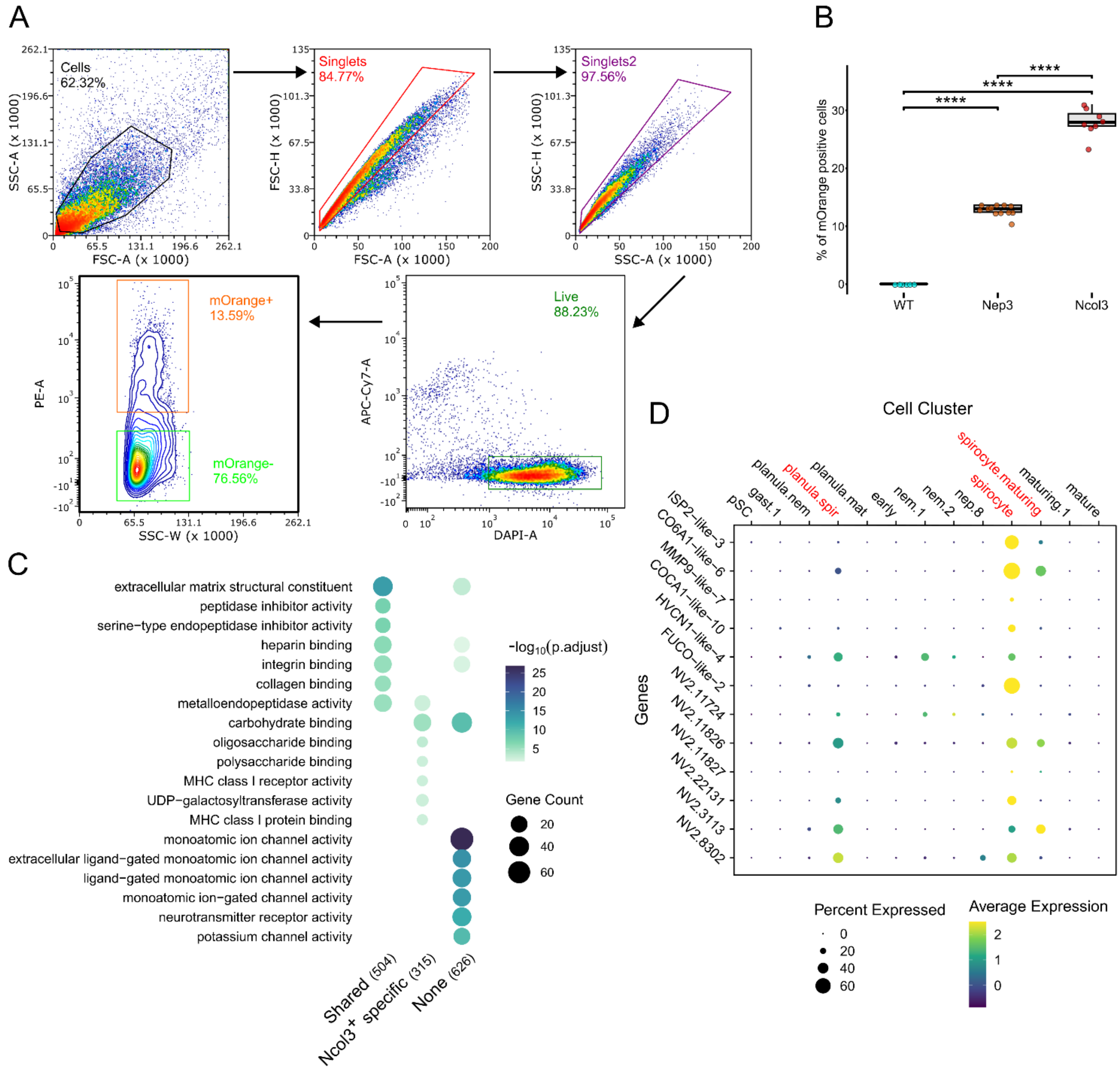


**Figure S2. FACS isolation and transcriptional profiling of cnidocyte reporter lines. (A)** Flow cytometry gating strategy for isolation of mOrange-positive cells from transgenic animals. Sequential gates were applied to select cells, singlets, and live cells (Calcein Violet$^{+}$, Zombie NIR$^{-}$), followed by separation into mOrange$^{+}$ and mOrange$^{-}$ populations. Representative percentages for each gate are indicated. **(B)** Quantification of mOrange-positive cells across wild-type (WT), *Nep3::mOrange*, and *Ncol3::mOrange* animals. Statistical significance was assessed using one-way ANOVA followed by Tukey's HSD post hoc test (****$p < 0.0001$). **(C)** Gene Ontology (GO) enrichment analysis of differentially expressed genes for shared (Upregulated in Both Ncol3 and Nep3 positive cells), Ncol3$^{+}$ specific, and none. Circle size represents gene count and color indicates adjusted p-value. Dot plot showing expression of top anti-correlated genes (up in Ncol3$^{+}$, down in Nep3$^{+}$) across cnidocyte subclusters identified by single-cell RNA-seq (Cole *et al.* 2024). Dot size reflects the percentage of cells expressing the gene, and dot color represents average expression level.

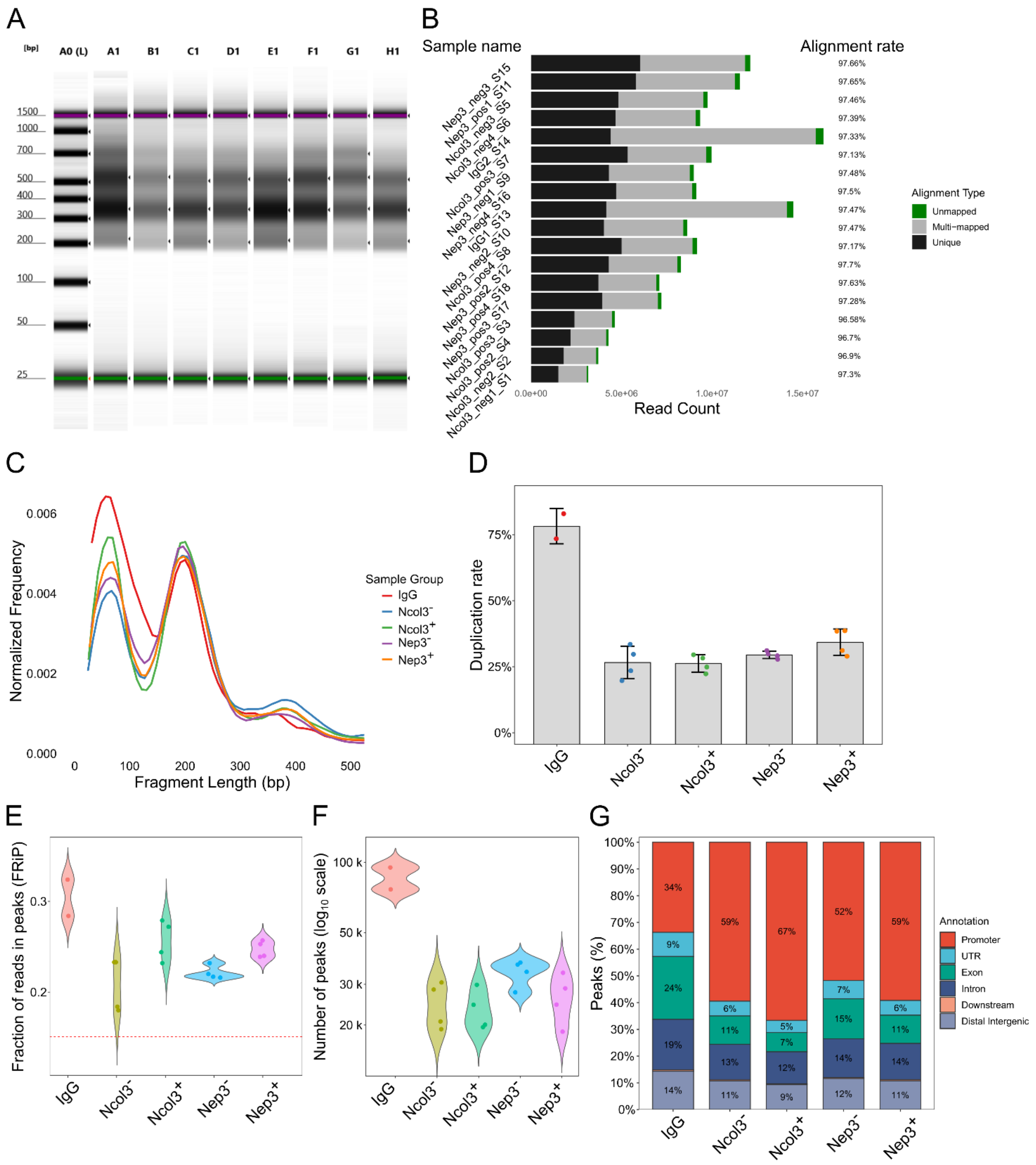


**Figure S3. Quality control and genomic characterization of CUT&Tag libraries. (A)** Bioanalyzer results showing fragment size distribution for representative CUT&Tag libraries across conditions. Prominent nucleosomal patterns are visible. **(B)** Read count and alignment statistics for all libraries. Bars show the proportion of uniquely mapped, multi-mapped, and unmapped reads. **(C)** Fragment length distributions of mapped reads, with clear mono- and di-nucleosome peaks detected across samples. **(D)** Duplication rates per library. IgG controls displayed higher duplication rates compared to Ncol3⁺, Ncol3⁻, Nep3⁺, and

Nep3⁻ samples. Bars represent mean ± SD**. (E)** Fraction of reads in peaks (FRiP) across samples. The red dashed line indicates a threshold of 0.15. **(F)** Number of peaks called per condition (log scale). **(G)** Genomic annotation of peaks across samples, showing the distribution across promoters, UTRs, exons, introns, downstream regions, and distal intergenic elements.

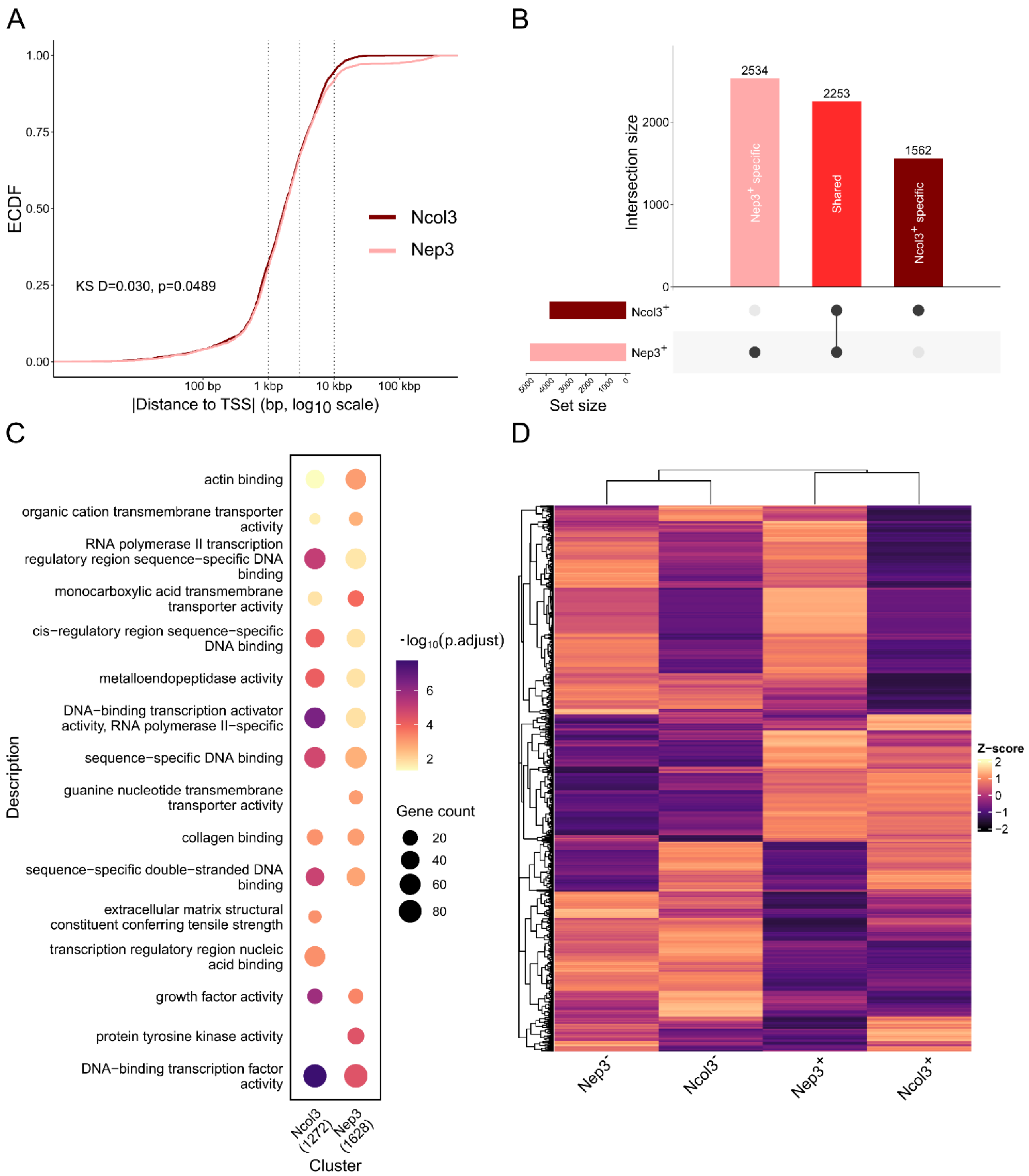


**Figure S4. Comparative analysis of Ncol3⁺ and Nep3⁺ enhancer landscapes. (A)** Empirical cumulative distribution function (ECDF) plot showing the distance of H3K27ac peaks to the nearest transcription start site (TSS) in Ncol3⁺ and Nep3⁺ cells. Peaks in both populations are preferentially located near TSSs, with a modest but significant difference between groups (Kolmogorov–Smirnov test). **(B)** UpSet plot showing the overlap of differential peaks between Ncol3⁺ and Nep3⁺ cells. Bars indicate the number of unique and shared peaks for each population. **(C)** Gene Ontology (GO) molecular function enrichment analysis of genes associated with population-specific peaks. Dot size represents the number of genes, and color indicates adjusted p-values. **(D)** Heatmap and hierarchical clustering of H3K27ac signal at differentially enriched distal peaks (> 3Kb) across conditions. Rows represent peaks, and columns represent aggregated samples of Ncol3$^{+}$, Ncol3$^{-}$, Nep3$^{+}$, and Nep3$^{-}$ populations. Z-scores of normalized signal intensity are shown.

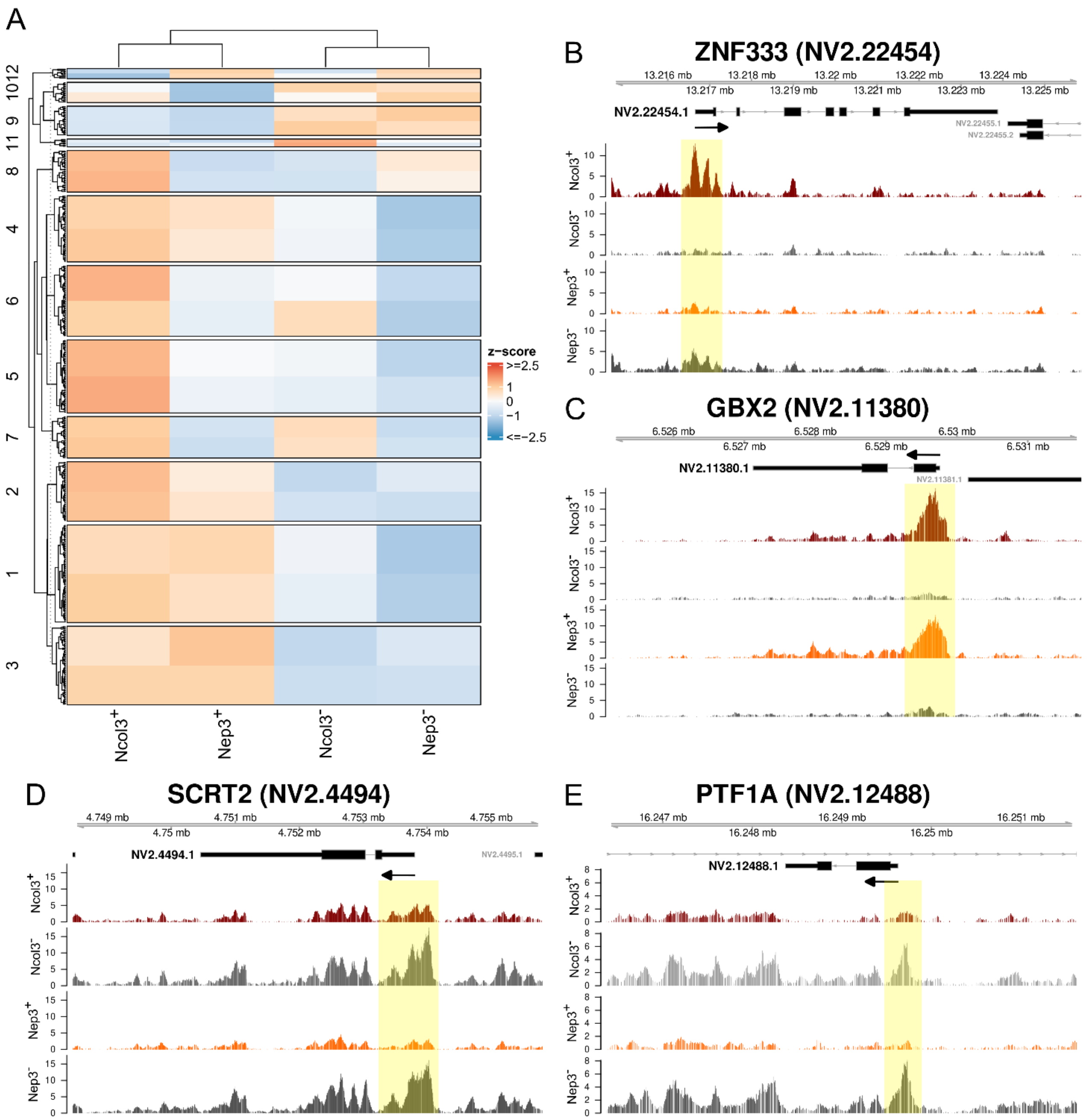


**Figure S5. Transcription factor associated H3K27ac regulation in cnidocyte cell types. (A)** Heatmap showing clustered patterns of H3K27ac enrichment at transcription factor associated peaks across Ncol3$^+$, Ncol3$^-$, Nep3$^+$, and Nep3$^-$ populations. Rows represent peak clusters, and columns represent sample groups. Signal intensity is displayed as Z-scores. **(B–E)** Genome browser views of representative loci showing cell type-specific activity near transcription factor genes. Highlighted regions (yellow) indicate enriched H3K27ac peaks. Examples include ZNF333 (NV2.22454) (B), GBX2 (NV2.11380) (C), SCRT2 (NV2.4494) (D), and PTF1A (NV2.12488) (E). Tracks display normalized CUT&Tag signal for each condition.

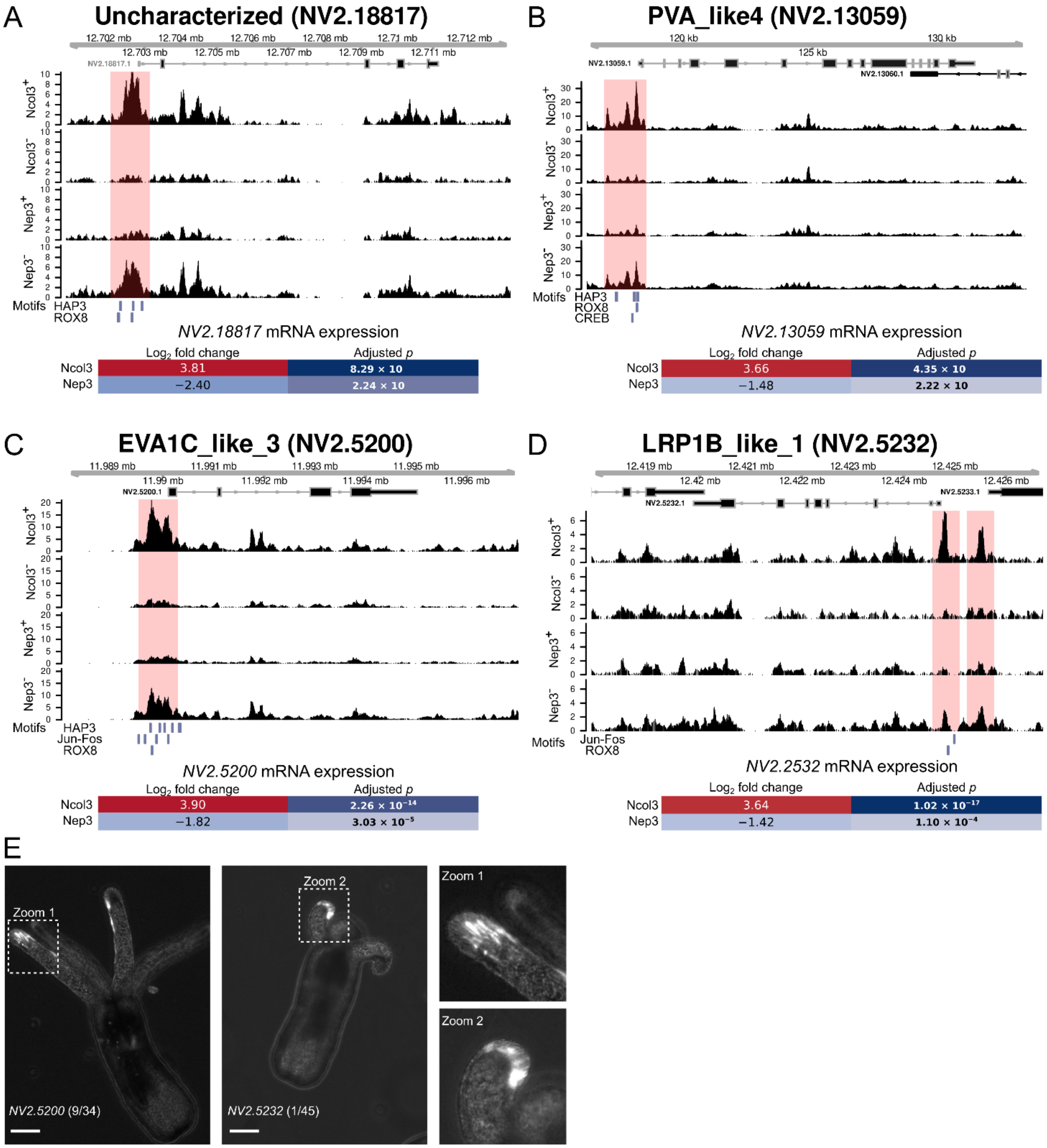


**Figure S6. Functional validation of candidate regulatory elements using transgenic reporter assays. (A-D)** Genome browser views of CUT&Tag H3K27ac signal at loci containing candidate enhancer regions. Highlighted areas (pink) mark enriched peaks tested in reporter constructs. Associated genes include an uncharacterized gene (NV2.18817) **(A)**, PVA_like4 (NV2.13059) **(B)**, EVA1C_like_3 (NV2.5200) **(C)**, and LRP1B_like_1 (NV2.5232) **(D)**. Motif analysis of these regions revealed putative transcription factor binding sites (bottom). Heatmaps below each track show RNA-seq expression changes ($log_2$ fold change and adjusted p-values) between Ncol3$^+$ and Nep3$^+$ cells.**(E)** Representative images of transgenic animals carrying reporter constructs driven by candidate regulatory regions upstream of mTurquoise2. Reporter expression was observed in cnidocytes of tentacles injected with the NV2.5200 (C) and NV2.5232 (D) constructs. Zoomed insets highlight expression patterns. Numbers indicate the fraction of positive animals out of inspected embryos. Scale bars: 100 μm.
